# Tissue-embedded CD4⁺ plasticity defines mucosal immunity in Inflammatory Bowel Disease

**DOI:** 10.1101/2025.10.31.685649

**Authors:** Qinyue Jiang, Veerle A. Merkus, Ciska Lindelauf, Nannan Guo, Laura F. Ouboter, Thomas Höllt, Philip W. Voorneveld, Caroline R. Meijer-Boekel, Frits Koning, Andrea E. van der Meulen-de Jong, Maria Fernanda Pascutti, Vincent van Unen

**Author notes:** **Corresponding author:** Vincent van Unen, **Address:** Postzone E3-Q, P.O. Box 9600, 2300 RC Leiden, The Netherlands.

## Abstract

CD4⁺ T helper (Th) cell responses to commensal microbiota are linked to Inflammatory Bowel Disease (IBD), yet how Th programs coexist and evolve in human tissues remains poorly defined. Here, we profiled CD4⁺ memory T cells in intestinal biopsies using immunological and histological approaches to map their phenotypes, functional states, and spatial relationships across disease states. A marked expansion of CD4⁺ T cells concomitant with a RORγt⁺ Th population with elevated T-bet expression was linked to progression of inflammation. Moreover, Foxp3⁺ cells co-expressing RORγt emerged within the inflamed niche, indicating regulatory–Th17 plasticity. Trajectory visualization revealed a potential branched differentiation path towards either regulatory or tissue-resident Th17-like fates, with both termini expressing activation and proliferation markers. Correlation network analysis connected pro-inflammatory CD4⁺ states to T-bet⁺Granzyme-B⁺ CD8⁺ subsets, indicative of crosstalk between helper and cytotoxic lineages. Histology corroborated this organization, showing frequent interactions between CD4⁺ and CD8⁺ cells in the lamina propria and epithelial border. In functional assays, TCR stimulation during active disease revealed broad suppression of CD4⁺ pro-inflammatory cytokines concomitant with expansion of Foxp3⁺ cells. Conversely, HLA-DR^+^CD38^+^ memory subset retained multifunctionality, producing elevated levels of pro-inflammatory cytokines. Together, these results provide insight into a dynamic, tissue-embedded CD4 landscape in IBD.

**Highlights:**

1. IBD inflammation shapes CD4^+^ T-cell plasticity and tissue-residency programming.
2. Correlated CD4^+^ and cytotoxic CD8^+^ T cells co-localize in inflamed mucosa.
3. Mucosal CD4⁺ memory T cells show hypo-responsiveness in active IBD.
4. HLA-DR^+^CD38^+^ memory CD4⁺ T cells amplify inflammation via cytokine output.

## Introduction

Disrupted immune responses to the gut microbiota in genetically susceptible individuals, potentially triggered by largely unknown environmental factors, contribute to the pathogenesis of Inflammatory Bowel Diseases (IBD)^1^. IBD typically follows a chronic, relapsing-remitting course, and comprises two main subtypes: Crohn’s disease (CD) and ulcerative colitis (UC). UC affects the colon and is characterized by continuous mucosal lesions, whereas CD can involve any segment of the gastrointestinal tract and frequently displays transmural inflammation. Current therapeutic goals are to induce and maintain clinical, biochemical, and durable endoscopic remission to minimize bowel damage and improve long-term outcomes^2^. Neither CD nor UC has a single phenotype; both are heterogeneous with respect to clinical presentation, treatment responses, and immune profiles^3^. Management of disease remains largely reactive, with treatment adjustments commonly driven by worsening of clinical parameters. Molecular and cellular characterization that accounts for inter-individual variability in tissue pathology could better guide clinical management and advance personalized therapy in IBD^4^. Dysregulation of CD4⁺ T cell subsets is widely implicated in chronic intestinal inflammation. In IBD, an imbalance between inflammation-promoting and immunoregulatory populations sustains mucosal damage and persistent immune activation, ultimately contributing to the development of chronic disease^5^. Upon antigen recognition, naïve CD4^+^ T cells can differentiate into distinct effector subsets, including pro-inflammatory Th1, Th2, and Th17, as well as a regulatory T (Treg) subset^6, 7^. In the intestine, regional tissue-resident memory CD4⁺ T cells (T_RM_) provide long-term local immunity to microbial antigens, and can be rapidly reactivated, undergo transient proliferation, and reacquire distinct T-helper phenotypes^8^. Recent studies have highlighted that polarized T-helper cells can adopt alternative phenotypes and functionalities during disease progression, accompanied by changes in signature transcription factors and lineage-defining cytokine profiles^9, 10^. While bidirectional Treg–Th17 plasticity has been described in IBD^10, 11^, the mechanisms governing this switching in human intestinal tissues across disease states remain poorly defined.

Although CD4⁺ T cells are major contributors to chronic intestinal inflammation, the intestinal immune landscape includes diverse cell types that interact with CD4⁺ T cells across the lamina propria and intraepithelial compartments to shape adaptive responses in IBD. Among these, CD8^+^ T cells, located in proximity to CD4⁺ T cells, are thought to be implicated in epithelial injury in IBD^12, 13^. We previously identified an inflammation-associated immune network enriched for activated HLA-DR^+^CD38^+^ effector memory CD4^+^ T cells in treatment-naïve IBD patients^14^. However, the phenotype, function, and trajectory of this population, particularly during disease remission, remain poorly defined.

To address these gaps, we performed immunological functional assays with spectral flow cytometry on freshly obtained intestinal biopsies and paired peripheral blood from patients with active IBD or in clinical remission, as well as non-IBD noninflamed controls. Here, we show that IBD harbors a dynamic, tissue-embedded CD4⁺ landscape with inflammatory expansion that recedes in remission, regulatory-Th17 plasticity (Foxp3–RORγt), and branched trajectories toward regulatory and tissue-resident fates. We also show coordinated CD4–CD8 networks and a multifunctional HLA-DR⁺CD38⁺ memory subset that persists despite global cytokine suppression, pointing to the development of rational therapeutic strategies to rebalance resident/regulatory programs while preserving protective memory.

## Results

### CD4^+^ T cells are enriched in the inflamed mucosa

To comprehensively assess intestinal and circulating T cells, together with other conventional and non-conventional lymphoid subsets, we applied a validated 37-plex full spectral flow cytometric panel^5^ to single-cell suspensions of endoscopic intestinal samples (n=68) and peripheral blood mononuclear cells (PBMCs) (n=32) from 27 IBD patients (CD, n=17; UC, n=10) and 5 controls (clinical metadata listed in **Supplementary Table 1**), yielding a dataset on 3.5 million live CD45^+^ cells (**Supplementary Fig. 1A**). Based on marker expression profiles, we delineated the intestinal T cell compartment and identified 7 major subsets: CD4^+^ T cells, conventional CD8αβ^+^ T cells, CD8αα^+^ T cells, CD8αβ^+^ NKT cells, double-negative (DN) T cells (CD8α^−^CD8β^−^CD4^−^), mucosal-associated invariant T (MAIT) cells, and γδ T cells (**Fig. 1A**). Next, samples were stratified by disease status (control, noninflamed, inflamed and remission), disease subgroup (control, CD, UC), and intestinal location of the analyzed samples (colon and ileum) (**Fig. 1B**). Cell-frequency analysis revealed pronounced heterogeneity in immune composition, with CD4^+^ T cell subsets predominating in the inflamed mucosa (**Fig. 1B**). Density plots of an opt-SNE analysis highlight differential presence of the CD4^+^ T cell subsets depending on disease status (**Fig. 1C and Supplementary Fig. 1B**). Consistently, the frequency of CD4^+^ T cells was significantly increased during inflammation and decreased in remission across disease types (CD, UC) and sites (colon, ileum) (**Fig. 1D**). In contrast, frequencies of other major lineages declined (conventional CD8αβ^+^ T, NKT and γδ T cells) or remained unchanged (CD8αα^+^ T, DN and MAIT cells) (**Supplementary Fig. 1C**) in the inflamed mucosa.

**Fig. 1.**
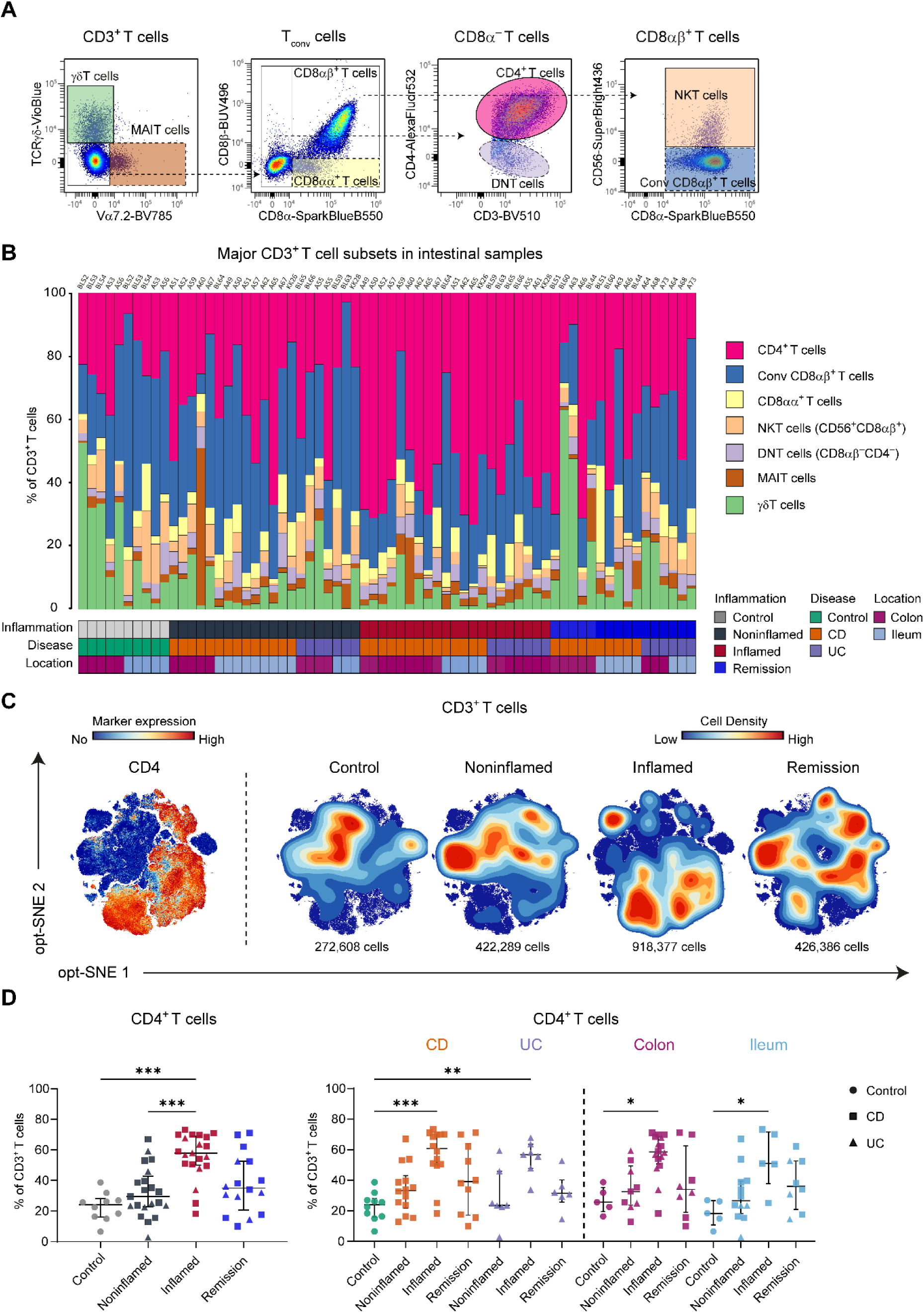
CD4⁺ T cells dominate in inflamed tissue samples in IBD. **(A)**, Bivariate plots showing the gating strategy to identify 7 major CD3⁺ T-cell subsets in intestinal samples, including CD4^+^ T, conventional CD8αβ^+^ T, CD8αα^+^ T, NKT (CD56^+^CD8αβ^+^), Double Negative (DN, CD8ab^−^CD4^−^), MAIT and γδ T cells. **(B)**, Bar graph showing cell frequencies of the 7 major CD3^+^ T-cell subsets. Depicted are the clinical metadata stratified for disease state(controls, noninflamed, inflamed, remission); disease subtype (control, CD and UC); and intestinal location (colon and ileum). **(C)**, opt-SNE embedding of the collective CD3^+^ T cells derived from 68 samples plots displaying CD4 expression in intestinal CD3⁺ T cells. Density maps showing the density of cells stratified for disease state. **(D)**, The frequencies of CD4⁺ T cells stratified for disease state, disease subtype and intestinal location, as percentage of total CD3⁺ T cells. Error bars indicate the median with interquartile range. *p≤0.05, **p≤0.01, ***p≤0.001, Kruskal–Wallis test with Dunn’s test for multiple comparisons.

### Inflammation shapes the CD4^+^ memory T cell landscape

Next, we performed a t-SNE analysis of intestinal memory T cells (T_M_), including the CD45RA^+^CCR7^−^, CD45RA^−^CCR7^−^, and CD45RA^−^CCR7^+^ subsets gated within the CD4^+^ population. Based on the marker expression profiles, hierarchical clustering of the collective 65,732 cells yielded 46 distinct T_M_-helper clusters (**Fig. 2A–B and Supplementary Fig. 2A**). The associated density plots display diverse distribution patterns related to disease state (**Fig. 2B**). Pearson’s rank correlation analysis identified four main cellular networks (**Supplementary Fig. 2B**), of which two (**Fig. 2C**) correspond to the high-density regions of the noninflamed (bottom clusters) and inflamed (top right clusters) groups shown in Fig. 2B, respectively. By analyzing the frequencies of those clusters, we observed that CD4-15, CD4-12 and CD4-48 were underrepresented during the active phase of disease (**Fig. 2D**), while CD4-13, CD4-5, CD4-7, CD4-18, CD4-21 and CD4-4 were notably increased in the inflamed group and decreased following the resolution of inflammation (**Fig. 2E**). The remaining clusters showed no significant quantitative changes.

**Fig. 2.**
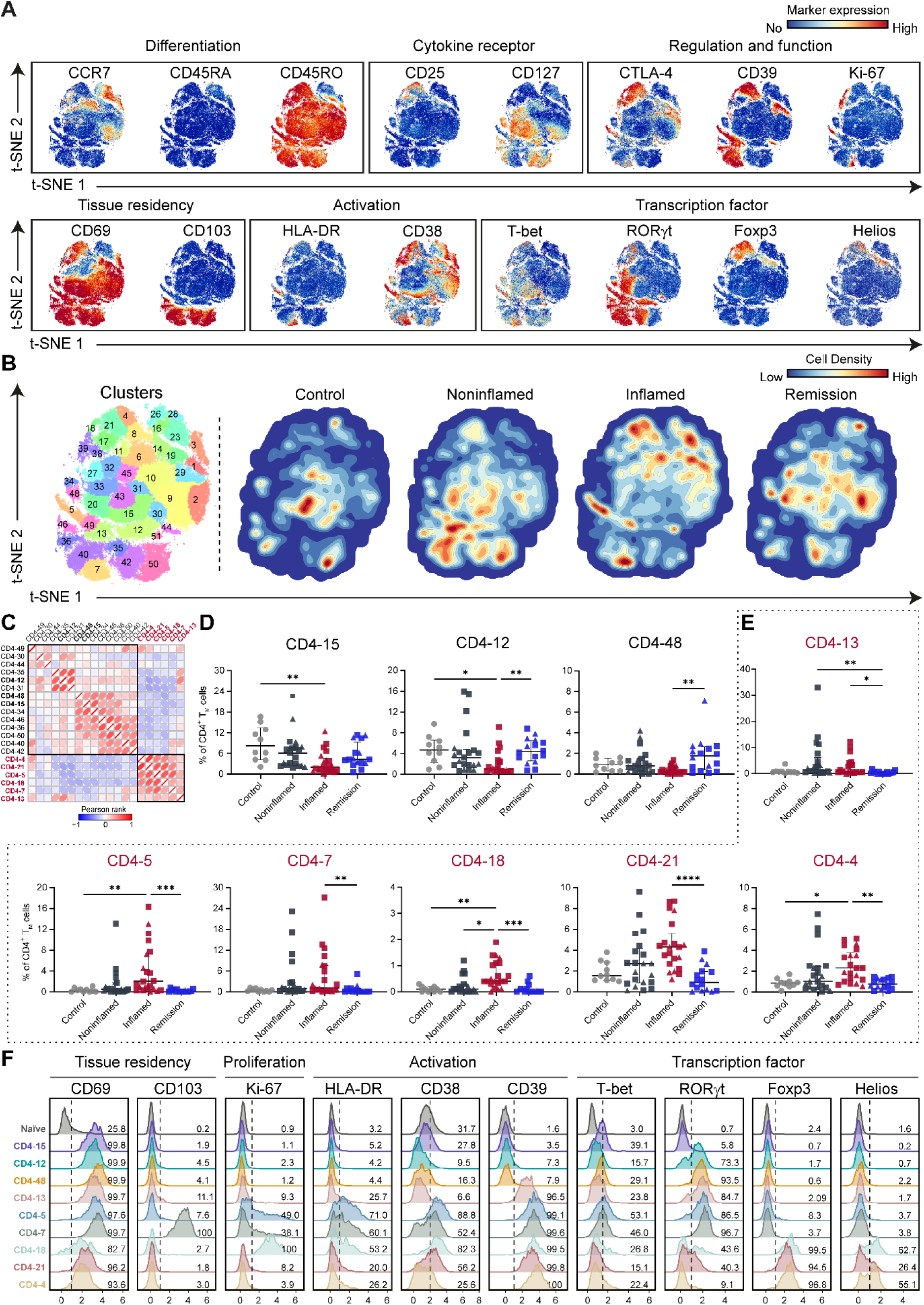
The Inflammation niche harbors several activated tissue-resident CD4 T cell subsets marked by distinct transcription factor expression profiles. **(A),** t-SNE plots embedding of 65,732 intestinal CD4⁺ memory T cells colored for the expression values of the indicated markers. **(B)**, t-SNE plot showing cluster partitions and density maps showing the abundance of cell subsets stratified for disease state. **(C)**, Pearson correlation matrix identifying a highly correlated cluster of CD4 T cell subsets (bottom right). **(D-E)**, Dot plots showing frequencies of indicated clusters stratified for disease state indicating that the CD4 T cell subsets in the correlated cluster identified in C are all increased in patients samples. Error bars indicate the median with interquartile range. *p≤0.05, **p≤0.01, ***p≤0.001, Kruskal–Wallis test with Dunn’s test for multiple comparisons. **(F)**, Overlay of histograms of CD4⁺ naive T cells (random selection of 1,000 naive cells from 68 samples) and IBD-associated clusters (derived from Fig. 2D**–E**) illustrating the expression patterns of markers related to tissue residency (CD69 and CD103), cell proliferation (Ki-67), activation (HLA-DR, CD38 and CD39) and transcription factors (T-bet, RORγt, Foxp3 and Helios).

To refine the phenotypic features of subsets involved in IBD inflammation, we overlaid marker expression across clusters (**Fig. 2F and Supplementary Fig. 2C**). Most CD4^+^ T_M_ clusters displayed CD45RA^−^CCR7^−^ effector-memory features, whereas a minority within CD4-18 and CD4-21 co-expressed low levels of CCR7, consistent with a central memory phenotype (**Supplementary Fig. 2C**). Compared to naïve T cells, all these CD4^+^ T_M_ subsets exhibited abundant expression of CD69, with cluster CD4-7 co-expressing CD103, indicating tissue-residency. Strikingly, the six inflammation-linked clusters (CD4-13, CD4-5, CD4-7, CD4-18, CD4-21, and CD4-4) co-expressed HLA-DR, CD38, and CD39 to varying degrees, consistent with activation. In addition, the expression of Ki-67 in CD4-5, CD4-7 and CD4-18 suggested proliferative potential. Distinct expression patterns of the transcription factors (T-bet, RORγt, Foxp3 and Helios) were observed among the CD4^+^ T_M_ clusters. For example, CD4-5 and CD4-7 co-expressed RORγt and T-bet, consistent with Th1/Th17-like states, but were otherwise phenotypically similar, differing primarily by CD103 expression restricted to CD4-7. Moreover, inflammation-associated CD4-18, CD4-21 and CD4-4 expressed Foxp3, with variable RORγt and Helios, indicating Treg identity and plasticity. Furthermore, we performed PCA on frequencies of 46 CD4^+^ T_M_ cell clusters (**Supplementary Fig. 2D**). Noninflamed samples (control, noninflamed IBD and remission) were clustered together and separately from inflamed samples. CD and UC samples were separated from the controls, with clear colon-ileum stratification. Overall, the inflammatory status and tissue location were the dominant source of variation.

Collectively, these findings suggest a disease-state–dependent, tissue-resident CD4⁺ memory landscape, with heterogeneous Th1/Th17 and Foxp3⁺ networks, whose frequencies stratify colon *versus* ileum and IBD *versus* control, increase with inflammation, and recede in remission.

### Trajectory analysis reveals bifurcating inflammatory paths of CD4⁺ memory T cell differentiation

Given the observed plasticity within CD4⁺ T-helper subsets, we next examined how these states evolve during inflammation. We used the dynamic t-SNE visualization in Cytosplore with naïve CD4⁺ T cells as a reference (**Fig. 3A and Supplementary Fig. 3A**)^15, 16^. Cells were initially unordered at onset of the t-SNE computation, but by stage 2 naïve cells segregated from the remaining memory cells. By stages 3–4 an interconnected network emerged with two branches bridged by cluster 13 (**Fig. 3A**). Disease-status overlays showed that the left branch comprised noninflamed clusters (CD4-12, CD4-15, CD4-48), whereas the right branch contained inflammation-linked clusters (CD4-5, CD4-7, CD4-4, CD4-18, CD4-21) (**Fig. 3B**). Marker expression profiles showed CD127 enrichment along the noninflamed branch and uniform CD39 positivity across inflamed clusters, with concurrent increases in CD38, HLA-DR and Ki-67 on the inflamed branch (**Fig. 3C**), consistent with activation and proliferation.

**Fig. 3.**
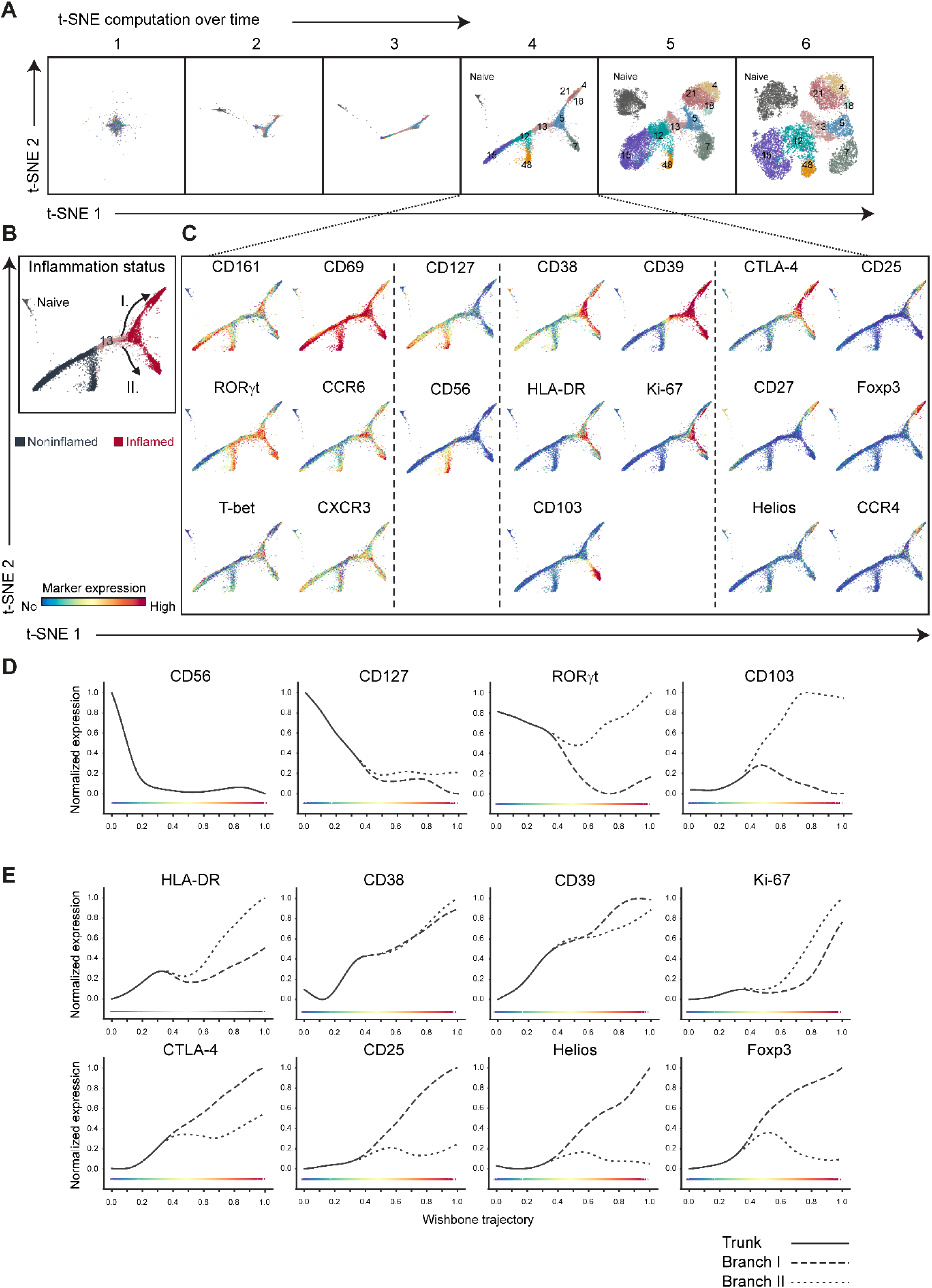
Potential differentiation trajectories of CD4 clusters during inflammation. **(A)**, t-SNE embeddings of CD4⁺ naive T cells (random selection of 1,000 naive cells from 68 samples) and the 9 CD4 clusters derived from Fig. 2D**–F**, showing single cells at six stages over the course of the t-SNE computation. Colors represent cluster partitions as described in Fig. 2F. **B**, stage 4 of the t-SNE embedding colored for inflammatory status and with arrows indicating 2 potential developmental trajectories associated with inflammation. **C**, Distinct marker expression profiles distinguishing the non-inflamed and inflamed tissue samples. **(D–E)**, Line plots generated by Wishbone (trajectory 0-1.0) showing the changes in normalized expression of indicated markers along bifurcating developmental trajectories.

To quantify marker dynamics along putative differentiation pathways, we applied Wishbone rooted at the intermediate cluster 13. This analysis revealed a clear bifurcation driven by divergent expression of RORγt/CD103 *versus* Foxp3/CTLA-4/CD25/Helios (**Fig. 3D–E**), delineating inflammatory trajectories toward a tissue-resident Th17-like fate (Branch I) and a regulatory fate (Branch II), respectively. Activation markers (HLA-DR, CD38, CD39, Ki-67) rose along the inflamed branch, supporting a model of inflammation-associated maturation from a shared intermediate. Additional overlays further corroborated branch identity (**Fig. 3C–E**): CD161 and CCR6, canonical Th17 features, were concentrated along the inflamed, tissue-resident branch (Branch I), whereas CCR4 and CD27, characteristic of Treg/less-differentiated memory, were biased toward the inflamed, regulatory branch (Branch II). CXCR3 aligned with T-bet–associated segments within Branch I, CD69 was broadly elevated across both branches, and a small CD56^+^ subset localized in the noninflamed branch, suggesting NK-like attributes.

Taken together, these analyses reveal a bifurcation from a shared intermediate into two inflammatory trajectories, one toward a tissue-resident Th17-like effector state and the other toward a regulatory fate, signifying inflammation-driven diversification of CD4^+^ T-helper memory programs.

### Coordination of CD4⁺ and CD8⁺ T-cell programs at the mucosal interface in IBD

To interrogate potential CD4–CD8 crosstalk in the intestine, we next characterized the CD8⁺ T-cell compartment. Unsupervised dimensionality reduction identified 20 phenotypically distinct CD8⁺ clusters (**Supplementary Fig. 4A**). We then integrated these CD8⁺ clusters with the CD4⁺ memory clusters and performed a Pearson correlation network analysis, resolving five multi-lineage cellular networks (**Fig. 4A**), characterized by strong within-network positive correlations and between-network anticorrelations. Network 3 contained the five inflammation-associated CD4⁺ clusters (CD4-4, CD4-5, CD4-7, CD4-18, CD4-21) together with five CD8⁺ clusters (CD8-2, CD8-3, CD8-8, CD8-9, CD8-10). Four of the five CD8⁺ clusters displayed a CCR7⁻CD45RO⁺CD45RA⁻ effector-memory phenotype, whereas CD8-3 was CCR7⁻CD45RO⁻CD45RA⁺, consistent with a terminally differentiated (T_EMRA_-like) state (**Fig. 4B and Supplementary Fig. 4B**). CD8-3, CD8-8, and CD8-9 showed a conventional CD8αβ phenotype; clusters 8 and 9 expressed Granzyme-B, indicating cytolytic potential, and CD8-9 additionally expressed CD103, consistent with tissue residency. In contrast, CD8-2 and CD8-10 lacked CD8β and co-expressed CD4 with CD8α, alongside RORγt and T-bet (**Fig. 4C and Supplementary Fig. 4B**). Notably, all five CD8⁺ clusters were significantly increased, or trended upward, in inflamed samples in parallel with their correlated CD4⁺ memory clusters (**Fig. 4D**). Conversely, four CD8⁺ clusters were reduced in inflammation (**Supplementary Fig. 4C**), indicating contraction of noninflamed/homeostatic states outside Network 3.

**Fig. 4.**
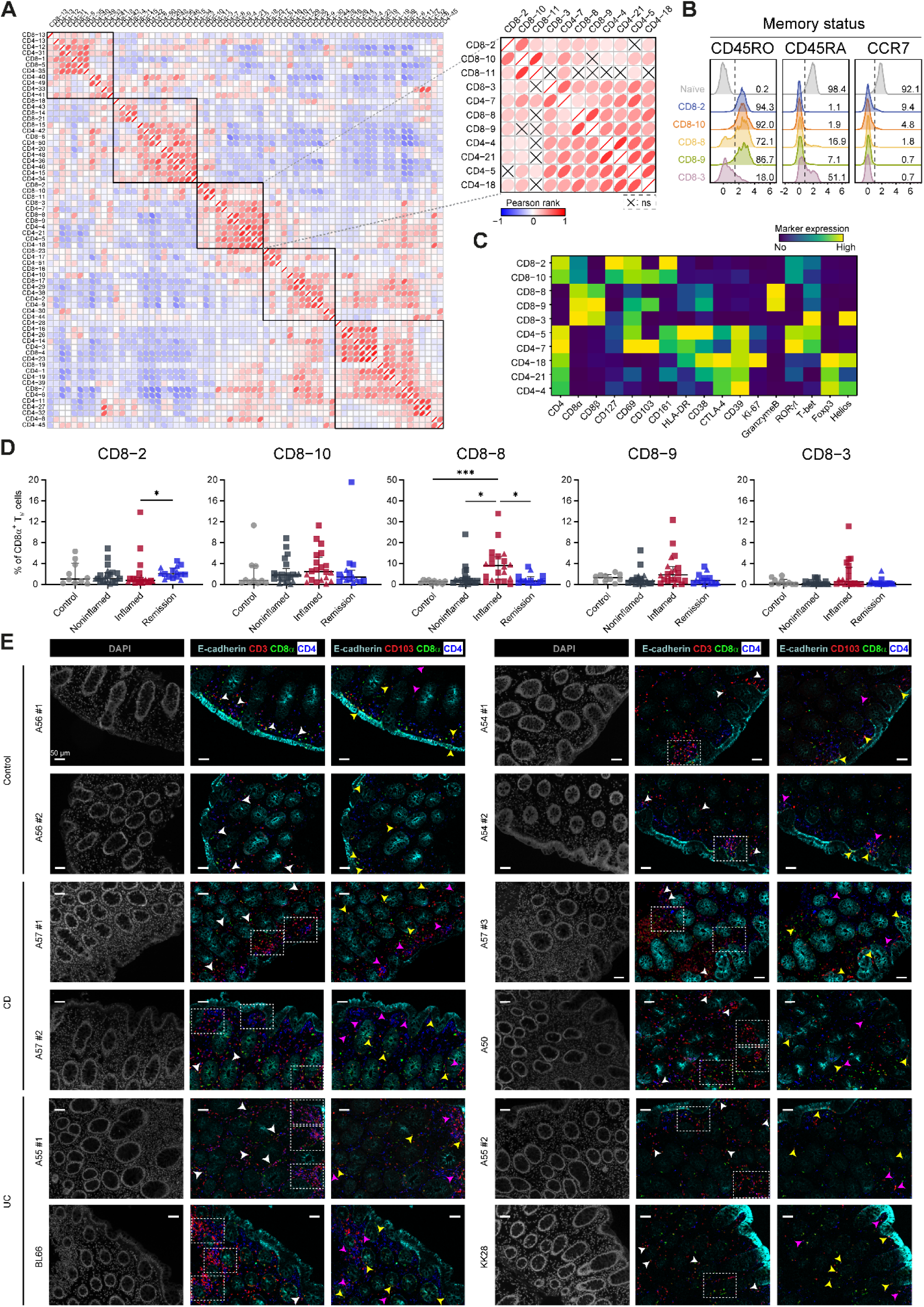
CD4–CD8 T cell crosstalk in the lamina propria. **(A)**, Pearson rank-based correlation matrix showing pairwise relationships among all memory CD4 clusters and CD8 clusters based on cell percentages (of total memory CD4⁺ and CD8α⁺ T cells) and a magnified view of the third network containing the five inflammation-associated CD4⁺ clusters together with five CD8⁺ clusters along with a statistical evaluation of inter-cluster correlations. “ns” denotes non-significant correlations. **(B)**, Overlay of histograms showing the CD45RO, CD45RA and CCR7 expression profiles on the correlated CD8 subsets and an unrelated naïve cluster for comparison. **(C)**, Heatmap showing median marker expression values of the correlated CD4 and CD8 clusters identified in **(A)**. **(D)**, The frequencies of CD8 clusters stratified for disease state, as percentage of total CD8α^+^ T cells. Error bars indicate the median with interquartile range. *p≤0.05, **p≤0.01, ***p≤0.001, Kruskal–Wallis test with Dunn’s test for multiple comparisons. **(E)**, Representative images of the detection of CD103 positive CD4⁺ and CD8α⁺ T cells in control, CD and UC samples, showing E-cadherin (colored in cyan), CD3 (colored in red), CD8α (colored in green), CD4 (colored in blue), and DAPI (colored in grey) as nuclear counterstain. Regions of CD4–CD8 cell colocalization within each ROI are marked by white arrows. Yellow arrows: CD3⁺CD8α⁺CD103⁺ cells; magenta arrows: CD3⁺CD4⁺CD103⁺ cells. Scale bar, 50 μm.

To define the spatial organization of resident CD4⁺ and CD8⁺ T cells in the inflammatory microenvironment, we performed six-color multispectral immunofluorescence on inflamed colonic tissue from patients with CD (n=2) and UC (n=3), as well as on noninflamed colonic tissue from healthy donors (n=2), displaying four representative regions of interest (ROIs) per group. We simultaneously detected E-cadherin, CD3, CD4, CD8, CD103, and DAPI (**Fig. 4E**). Compared with controls, IBD samples showed increased T-cell infiltration within the lamina propria and along the epithelial interface, with frequent juxtaposition of CD4⁺ and CD8⁺ cells (white arrows). CD103⁺ intraepithelial cells were predominantly CD8⁺ (yellow arrows), whereas CD103⁺CD4⁺ cells localized near the epithelial border or within the lamina propria (magenta arrows), consistent with coordinated resident programs at the mucosal interface. Notably, this spatial pattern accords with the trajectory analysis (**Fig. 3**), which revealed progressive CD103 induction along the inflammatory CD4⁺ branch, linking phenotypic residency programming to tissue positioning.

### Inflammation dampens TCR-mediated activation of CD4⁺ memory T cells

To assess CD4^+^ T cell function, we stimulated single-cell suspensions from intestinal biopsies with α-CD3/α-CD28 for 6 hours and profiled CD40L together with cytokines characteristic of Th1 (IL-2, TNFα, IFNγ), Th2 (IL-13, IL-4), Th17 (IL-17A, IL-21, IL-22), and Treg (IL-10) by spectral flow cytometry (**Supplementary Fig. 5A**). Data from noninflamed samples (controls, noninflamed IBD, remission) were pooled and compared with inflamed samples. Two experimental batches showed concordant trends and are presented separately in Supplementary Fig. 5B–C, and the combined cohort is shown in Figure 5.

**Fig. 5.**
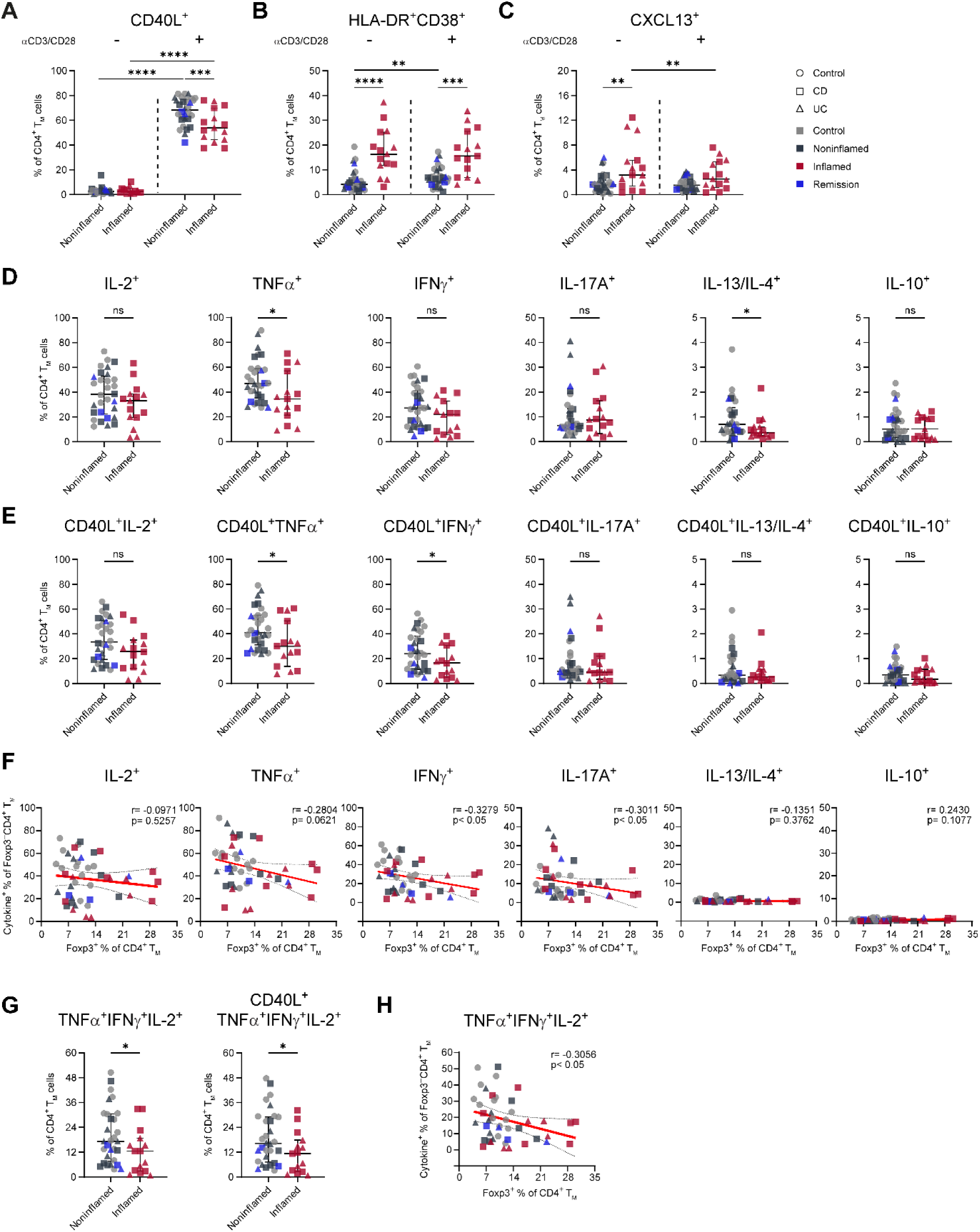
Functional profiling of intestinal CD4^+^ memory T cells. Intestinal suspensions were treated with α-CD3 and α-CD28 for 6 hours. Flow cytometry was used to quantify marker and cytokine expression. **(A–C)** The frequencies of CD40L⁺, HLA-DR⁺CD38⁺, and CXCL13⁺ cells in CD4⁺ memory T cells before and after stimulation from noninflamed and inflamed samples. **(D–E),** Intracellular expression of IL-2, TNFα, IFNγ, IL-17A, IL-13/IL-4 and IL-10 in total CD4⁺ memory T cells **(D)** and the responsive (CD40L⁺) memory T cells **(E)**. **(F)**, Correlations between frequencies of Foxp3⁻ CD4⁺ memory T cells producing IL-2, TNFα, IFNγ, IL-17A, IL-13/IL-4, IL-10 and frequencies of Foxp3⁺ CD4⁺ memory T cells were assessed using simple linear regression. Spearman’s rank correlation was used to determine correlation coefficients and *p* values. **(G),** The frequencies of TNFα⁺IFNγ⁺IL-2⁺ cells in CD4⁺ memory T cells and the responsive (CD40L⁺) memory T cells. **(H),** Correlation analysis showing the relationship between the frequencies of Foxp3⁻ CD4⁺ memory T cells producing TNFα, IFNγ and IL-2 simultaneously and Foxp3⁺ CD4⁺ memory T cells. Error bars indicate the median with interquartile range. “ns” denotes not significant, *p≤0.05, **p≤0.01, ***p≤0.001, Mann-Whitney test.

CD40L was robustly induced by stimulation in all groups but rose less in inflamed tissue, indicating attenuated activation (**Fig. 5A**). HLA-DR^+^CD38^+^ CD4⁺ T_M_ cells were higher at baseline in inflamed mucosa and remained at similar frequencies after stimulation (**Fig. 5B**). Although CXCL13⁺CD4⁺ T_M_ cells, indicative of a T follicular helper-like subset, were also increased at baseline in inflamed tissue, their frequencies were instead reduced after stimulation compared with noninflamed samples (**Fig. 5C**).

We next quantified cytokine production within the memory compartment (**Fig. 5D–E and Supplementary Fig. 5C**). After 6 hours, Th1 cytokines (IL-2, TNFα, IFNγ) predominated, Th17 cytokines (IL-17A, IL-21, IL-22) were lower in abundance, and Th2/Treg cytokines (IL-13, IL-4, IL-10) were largely rare. Moreover, TNFα was significantly decreased in the inflamed group, with IL-2 and IFNγ showing only mild reductions (**Fig. 5D**); these differences persisted when gating on activated CD40L^+^ cells (**Fig. 5E**). IL-17A and IL-21 were largely unchanged, while IL-22 trended lower (**Supplementary Fig. 5C**). IL-13/IL-4 showed a small but statistically significant difference with low absolute frequencies, and IL-10 showed no clear change (**Fig. 5D**). In the CD8⁺ memory compartment, CD40L expression did not change after stimulation, but IL-2 production was significantly reduced in inflamed versus noninflamed tissue, whereas TNFα, IFNγ and IL-17A were comparable (**Supplementary Fig. 6A-B**).

To investigate mechanisms of reduced cytokine production, we related Foxp3⁻ effector output to Foxp3^+^ Treg abundance. Significant negative correlations with Foxp3^+^ frequency were observed for IFNγ and IL-17A, with a trend for TNFα (**Fig. 5F**). Th2 cytokines and IL-10 were not significantly correlated. In addition, a multifunctional subset co-producing IL-2, TNFα and IFNγ was likewise diminished in inflamed tissue upon stimulation and inversely associated with Foxp3^+^ frequency (**Fig. 5G–H**).

Applying the same stimulation to paired blood samples revealed comparable CD40L induction and cytokine production between inflamed and noninflamed groups (**Supplementary Fig. 6C-E**), suggesting that the hypo-responsiveness is localized to the tissue.

Collectively, despite intact proximal activation through TCR engagement, CD4⁺ T_M_ cells from inflamed mucosa display selectively blunted Th1 effector function and reduced multifunctionality, features that are tissue-localized and in part linked to increased regulatory burden.

### HLA-DR⁺CD38⁺ memory CD4⁺ T cells in IBD are tissue-resident, proliferative, and multifunctional

We previously noted an expansion of activated HLA-DR^+^CD38^+^ effector-memory CD4^+^ T cells in treatment-naïve IBD^14^. Here, including both active disease and remission, we confirmed that HLA-DR and CD38 were upregulated across central memory (T_CM_), effector memory (T_EM_) and terminally differentiated effector memory (T_EMRA_) subsets (**Fig. 6A**), and that the HLA-DR^+^CD38^+^ fraction was increased in inflamed tissue and reduced in remission (**Fig. 6B**).

**Fig. 6.**
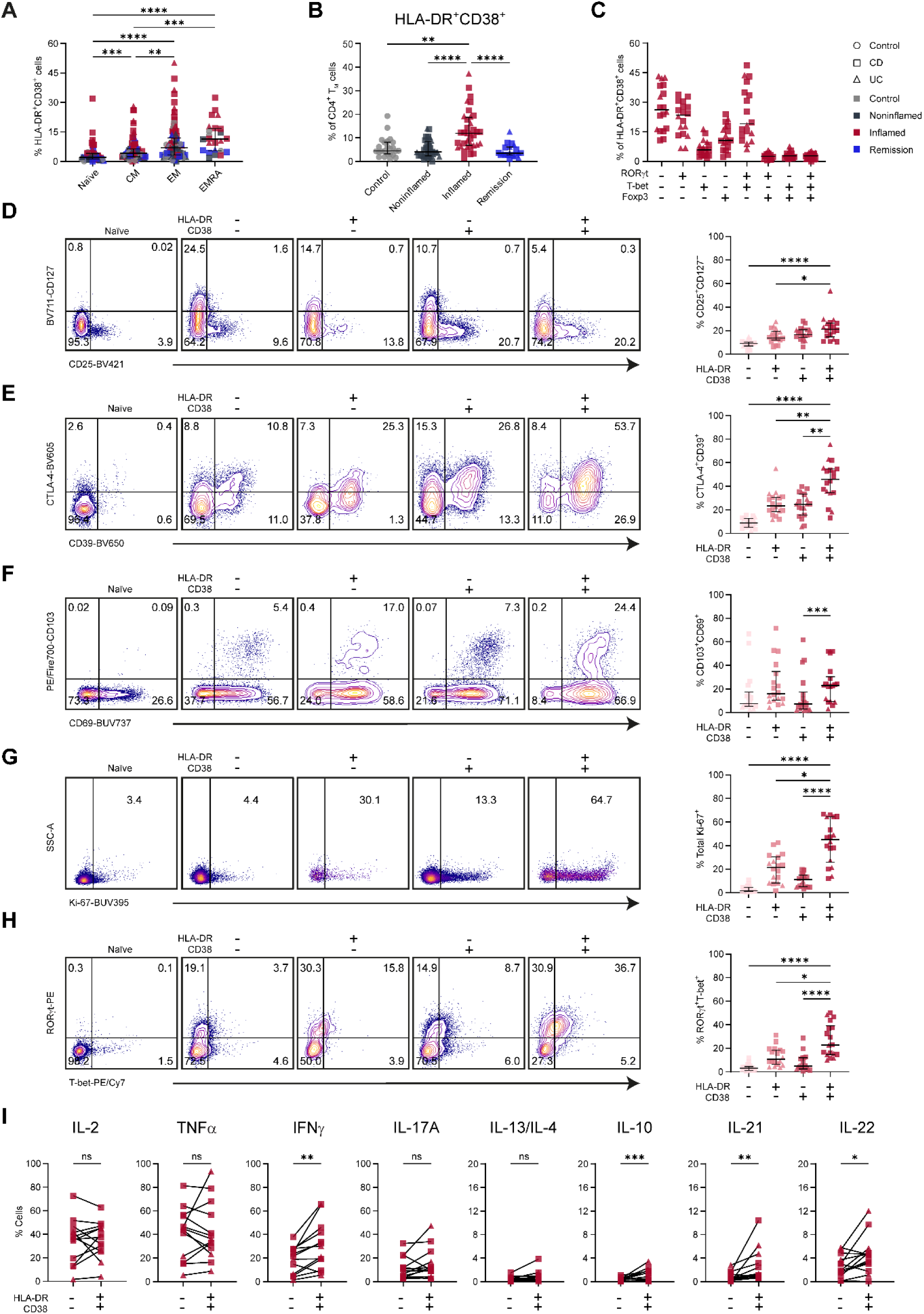
Phenotypic characterization and functional profile of IBD associated HLA-DR^+^CD38^+^ cells. **(A)**, Dot plots showing the percentage of double positive HLA-DR and CD38 cells in the naive, central memory (CM), effector memory (EM), and terminally differentiated effector memory (EMRA) CD4⁺ T cells**. (B)**, The frequencies of double positive HLA-DR and CD38 cells stratified for disease state, as percentage of total CD4⁺ memory T cells. **(C)**, Distribution of cells co-expressing various combinations of the three transcription factors (RORγ, T-bet and Foxp3) among HLA-DR⁺ CD38^+^ CD4^+^ memory T cells. **(D–H)**, The frequencies of CD25⁺CD127⁺, CTLA-4⁺CD39⁺, CD103⁺CD69⁺, Ki-67⁺ and RORγ⁺T-bet⁺ cells compared among four groups defined by HLA-DR (positive or negative) and CD38 (positive or negative) expression, shown in biaxial plots (left, naïve cells are shown for comparison) and dot plots (right) (data from one representative experiment).**(I)**, Comparison of the frequencies of IL-2, TNFα, IFNγ, IL-17A, IL-13/IL-4 and IL-10 positive cells in HLA-DR⁻CD38⁻ and HLA-DR⁺CD38⁺ populations. Error bars indicate the median with interquartile range. *p≤0.05, **p≤0.01, ***p≤0.001, “ns” denotes not significant, Kruskal–Wallis test with Dunn’s test for multiple comparisons or Wilcoxon tests, as appropriate.

Analysis of the transcription factor composition demonstrates that within HLA-DR^+^CD38^+^ memory cells, a large subset (∼25%) was negative for the lineage factors T-bet, RORγt and Foxp3, with single-positive Foxp3^+^ cells comprising a minority (∼10%) (**Fig. 6C**). Nonetheless, RORγt expression was clearly enriched (∼23%) and included a subset co-expressing T-bet (∼20%). Cells co-expressing Foxp3 and Helios, markers linked to Treg stability, were present at lower but increased levels in HLA-DR⁺CD38⁺ cells compared with HLA-DR⁻CD38⁻ cells (**Supplementary Fig. 7A–C**), indicating a small embedded regulatory-like subset.

Next, we compared the phenotypic profiles among HLA-DR/CD38-defined memory subsets within inflamed tissue. Compared with HLA-DR⁻CD38⁻ memory cells, the HLA-DR⁺CD38⁺ population showed higher CD25 and lower CD127 (**Fig. 6D**), indicating enhanced IL-2 receptor signaling and recent activation. They also upregulated CTLA-4 and CD39 (**Fig. 6E**), consistent with engagement of regulatory pathways and ectonucleotidase activity that can generate immunomodulatory adenosine. The same cells expressed increased CD103 and CD69 (**Fig. 6F**), markers of tissue residency and epithelial retention, and exhibited elevated Ki-67 (**Fig. 6G**), indicating ongoing proliferation. T-bet and RORγt were enriched in the HLA-DR⁺CD38⁺ subset (**Fig. 6H**), supporting polarization toward Th1/Th17 programs. Cells single-positive for HLA-DR or CD38 generally displayed intermediate levels across these features (**Fig. 6D–H**), placing them between quiescent double-negative and fully activated double-positive states.

Finally, we assessed cytokine production by HLA-DR^+^CD38^+^ CD4⁺ T_M_ cells. Upon stimulation with α-CD3/α-CD28, this subset produced more IFNγ, IL-21, IL-22, IL-10 than in HLA-DR⁻CD38⁻ counterparts (**Fig. 6I**). In contrast, IL-2, TNFα, IL-17A and Th2 cytokines (IL-13/IL-4) were comparable between groups (**Fig. 6I**), pointing to selective multifunctionality. t-SNE visualization of cytokine expression within the HLA-DR⁺CD38⁺ compartment (**Supplementary Fig. 7)** showed that activated Th1, Th17, Tfh/Th22-like cells largely occupied distinct regions with only limited overlap; within this landscape, polyfunctional IFNγ⁺IL-2⁺TNFα⁺ Th1-type cells predominated, while IL-17A was mainly produced by separate cells, with only a small subset co-expressing IL-17A together with these Th1 cytokines. Taken together with their activation, residency, and proliferative features, these data position HLA-DR⁺CD38⁺ CD4⁺ T_M_ cells as tissue-embedded effectors that can both amplify inflammation and modulate it through targeted cytokine programs.

## Discussion

Our findings show that inflammation in IBD reshapes the intestinal T cell landscape, marked by an expansion of activated CD4^+^ memory T cells and contraction of other T cell subsets. These CD4^+^ memory T cells displayed a tissue-resident, proliferative phenotype and correlated with specific cytotoxic CD8⁺ T cell subsets, suggesting coordinated mucosal immune responses. Their numbers declined in remission, underscoring inflammation-dependent expansion. We found these cells to be functionally hyporesponsive upon TCR stimulation, producing reduced TNFα and IFNγ and displaying limited polyfunctionality. This functional restraint was inversely correlated with local Foxp3^+^ Treg abundance. However, a specific CD4^+^ memory T cell subset expressing HLA-DR and CD38 maintained polyfunctionality in inflamed IBD. In essence, active IBD lesions are characterized by the coexistence of a broadly hyporesponsive CD4⁺ memory pool and a smaller HLA-DR⁺CD38⁺ subset that remains highly multifunctional, suggesting differential regulation within the wider CD4⁺ compartment under chronic inflammatory conditions.

Our high-dimensional approach also captured several ‘unconventional’ T-cell subsets (γδ T cells, MAIT cells and NKT-like cells)^17, 18^. We noted that γδ T cells, which normally reside abundantly in intestinal epithelium, were markedly decreased in the inflamed mucosa in our IBD cohort. This aligns with reports that mucosal γδ T cells are perturbed in IBD, although findings have varied^19^. Similarly, invariant NKT cells (CD1d-restricted T cells) were diminished in our inflamed samples. Consistently, clinical studies have reported that type 1 NKT cells are reduced in the blood and intestine of CD and UC patients, whereas type 2 NKT cells can accumulate in UC lamina propria, potentially contributing to typical Th2 responses^20^. MAIT cells typically accumulate in the gut and help regulate responses to microbiota^21–23^; however, we did not observe significant changes in their frequency in our dataset, in contrast to previous studies. Nonetheless, our observations reinforce the idea that intestinal inflammation can differentially affect innate-like T-cell populations.

A central finding of our study is the plasticity of CD4^+^ T-helper cells under inflammatory conditions. We identified RORγt^+^ Th17-lineage subsets with divergence into Th1-like (T-bet^+^IFNγ^+^) or Treg-like (Foxp3^+^CTLA-4^+^) fates, indicating context-dependent bifurcation within the IBD mucosa. In our cohort, mixed Th1/Th17 subsets (clusters CD4-13, CD4-5, and CD4-7) were selectively expanded in patients with active disease and returned toward baseline frequencies in patients in remission, directly linking these plastic states to clinical inflammation. This supports the concept that Th17 cells are highly flexible, adopting pro- or anti-inflammatory phenotypes depending on the local cytokine milieu^24, 25^. In line with this, HLA-DR⁺CD38⁺ EM CD4⁺ T cells, previously also identified as a central disease-associated population in treatment-naïve IBD^14^ and as activated Th1/Th17 cells in CD-associated perianal fistulas^26^, likely represent a clinically relevant manifestation of this plastic Th17/Th1 continuum. Some studies have shown that IL-23 promotes the emergence of pathogenic Th17 cells that gain IFNγ-producing, Th1-like features, including IL-17A⁺IFNγ⁺ and ex-Th17 effectors, particularly in T cell–transfer colitis models^27–29^. Blocking IL-23 in these models attenuates these pathogenic populations and ameliorates colitis, and, in patients, antibodies targeting IL-12/23p40 or IL-23p19 induce and maintain remission in both CD and UC^30–33^. Moreover, Heredia *et al.* demonstrated that IL-12 can drive gut-homing TIGIT⁻CD38⁺ memory T cells towards an ex-Th17/Th1-like pathogenic phenotype in a subset of pediatric CD with a severe disease course^34^. In contrast, IL-17 neutralization failed to show efficacy in IBD and in some trials was associated with worsening disease and paradoxical intestinal inflammation, likely due to its role in mucosal integrity. These findings underscore the need to recalibrate, not eliminate, Th17 responses.

Our trajectory analysis further supports a model in which Th17-like cells can shift toward Foxp3^+^ regulatory phenotypes in response to inflammation. We observed RORγt⁺Foxp3^+^ double-positive cells enriched in inflamed tissue, potentially representing Th17-derived Tregs. Expression of Foxp3 in RORγt⁺ cells may in part reflect activation rather than bona fide regulatory function; however, these cells also upregulated CD39 and CTLA-4, consistent with a Treg-like program potentially adapted to suppress Th17-skewed responses. Previous studies similarly suggest that Tregs are also plastic and can transiently adopt lineage-defining transcription factors to better control specific effector responses^35–37^. Interestingly, we noticed these cells simultaneously expressing CD161, though it is typically linked to a RORγt-driven transcriptional program and denotes a pro-inflammatory fate. This observation is consistent with previous findings that reported existence of CD161⁺ regulatory T cells with preserved suppressive capacity^38^. Conversely, Foxp3^+^ Tregs may become unstable under inflammatory pressure^39^, losing Foxp3 and acquiring effector functions. In support of this, single-cell profiling of ileal CD4⁺ T cells in CD identified proinflammatory FOXP3⁺ Treg subsets with TNFα-driven gene programs, impaired suppressive function, and enhanced cytokine production^40^. Thus, bidirectional plasticity between Th17 and Treg lineages may dynamically tune inflammation and resolution. Therapeutically, this suggests restoring the Th17/Treg balance as an attractive target. Low-dose IL-2, which preferentially expands Tregs, has shown safety and partial efficacy in UC^41^, though Treg instability remains a concern. Combining Treg-expanding strategies with cytokine blockade (e.g., IL-6/STAT3 inhibition) or next-generation antigen-specific or CAR-engineered Tregs may improve durability^39, 42, 43^.

Our data also highlight that clusters on the Th17-like inflammatory trajectory upregulated CD69 and CD103, indicating a transition to tissue-resident memory T cells. Tissue-resident T cells reside in the gut mucosa for decades^44^, where they can react swiftly to local stimuli^45^. In line with this, CD4⁺ T_RM_ have been shown to be expanded and to constitute a major TNFα-producing population in active CD^46^, supporting their pathogenic potential. Recent work in CD identified pathogenic CD103⁺CD161⁺CCR5⁺ CD4⁺ T cells with innate-like IFNγ responses and epithelial localization^47^. These cells exhibited innate-like effector functions, produced large amounts of IFNγ upon stimulation, and clustered near the epithelium. We similarly observed inflammation-linked CD4⁺ tissue-resident memory cells (exemplified by cluster 7), which likely represent the long-lived pathogenic effectors that embed in the gut tissue and could drive disease recurrence or persistence. The presence of such inflammation-prone resident memory T cells might explain why some patients relapse quickly after treatment withdrawal.

Correlation network analysis of our data demonstrated that the inflammation-associated CD4⁺ clusters were tightly linked to certain CD8⁺ T-cell clusters (forming a multi-lineage network). These CD8 subsets, which expanded in active disease, included CD8αβ⁺ cytotoxic tissue-resident T cells and unconventional CD4^+^CD8α^+^CD8β^−^ tissue-resident T cells, previously described by Lockhart et al^48^, co-expressing T-bet and RORγt in our data. Moreover, our findings are consistent with reports that ileal CD8⁺ tissue-resident memory T cells acquire context-dependent inflammatory programs in IBD^49^. Jaeger *et al.* further reported a spatial redistribution of CD8⁺ T cells in CD, with a loss of CD8⁺ IELs and a concomitant increase in CD103⁺ resident cells within inflamed tissue, suggesting that the depletion of CD8⁺ IELs may weaken epithelial barrier defense, while their accumulation in the lamina propria, together with CD4⁺ T cells, could amplify inflammation^50^. Spatially, multiplex immunofluorescence revealed dense aggregates of CD4⁺ and CD8⁺ T cells intermingled in inflamed IBD tissues. CD8⁺ T cells (especially CD8αβ⁺CD103⁺ IELs) were observed within or just below the epithelium, often adjacent to CD4⁺ cells, pointing to possible crosstalk. This coordinated localization suggests a scenario where CD4⁺ T cells and CD8⁺ cytotoxic T cells jointly contribute to epithelial injury in IBD.

Our study has limitations that open avenues for further research. Our data are cross-sectional; longitudinal sampling before, during, and after therapy would be valuable to observe T-cell state transitions and confirm which changes are causes or consequences of healing. Technical aspects like the use of *ex vivo* stimulation should be expanded with antigen-specific stimulation assays to mimic chronic exposure to microbial antigens. The duration of T-cell activation should be extended to elicit a broader range of cytokines, and provide clearer evidence of functional exhaustion. Finally, it would be intriguing to molecularly characterize the epigenetic landscape of these inflammation-adapted T cells.

In summary, this study provides an integrated view of how inflammation sculpts the intestinal T-cell landscape in IBD, revealing distinct T-cell states linked to disease activity. We propose that IBD flares are driven by a network of tissue-resident, Th17/Th1-skewed CD4⁺ memory T cells and cytotoxic CD8⁺ T cells positioned at the mucosal interface, whose pathogenic activity is counterbalanced by an expanding regulatory T-cell compartment. When inflammation subsides (either naturally or via treatment), this pathogenic network contracts and the regulatory arm regains dominance, restoring immune homeostasis. These findings underscore the plasticity of mucosal T cells in inflammation, likely shaping patient-specific outcomes. Targeting pathogenic T-cell memory while reinforcing regulatory circuits may offer a path to durable IBD remission, offering opportunities for the development of more targeted therapeutics.

## Materials and Methods

### Patients

IBD patients (minimum age 12 years old) were recruited from the Department of Gastroenterology and the pediatric department at Leiden University Medical Center (LUMC). Patients either had a prior diagnosis of IBD or underwent diagnostic ileocolonoscopy based on clinical suspicion of new-onset IBD. Controls were patients ≥ 18 years old known with another disease, such as history of colorectal cancer or familial colon cancer, who underwent a surveillance ileocolonoscopy. The studies were reviewed and approved by the Medical Ethical Committee of LUMC. After informed consent, fresh biopsies and peripheral blood were obtained directly at the endoscopy room according to protocol P15.193. Fresh samples were processed (see below) and results were stored in the LUMC-IBD biobank (protocol RP25.075).

The phenotype characterization cohort included 68 intestinal samples from 27 patients (CD, n=17; UC, n=10) and controls (n=5). T cell receptor stimulation assays were conducted with two cohorts of patients. The first included 20 intestinal samples from 6 IBD patients (CD, n=3; UC, n=3), and 3 controls, while the second included 25 intestinal samples from 10 patients (CD, n=5; UC, n=5) and 4 controls. Samples from 3 of these controls and 7 of these patients (CD, n=3; UC, n=4) were utilized for phenotypical characterization as well.

### Sample and clinical data collection

The clinical characteristics of all patients are shown in Supplementary Table 1. Samples from rectum, colon and ileum, both from inflamed and endoscopic noninflamed mucosa (if available) were collected from IBD patients. Noninflamed mucosa were bowel segments never inflamed or bowel segments currently in endoscopic remission (previously involved segment). Peripheral blood was drawn from all participants at the time of endoscopy.

Patients were classified according to the Montreal classification regarding age at diagnosis, location/extent of disease and behavior. In CD patients, the inflamed segment of each patient was classified according to the SES-CD, as inactive (0-2), mild (3-6), or moderate-severe (≥7). In UC patients, severity of disease was scored according to the endoscopic Mayo score, as inactive (0), mild (1) or moderate-severe (2-3).

### Isolation of single cells from endoscopic intestine and peripheral blood

Single-cell suspensions were prepared from fresh endoscopic biopsies as previously described^5^. Briefly, tissue samples were collected in Hank’s balanced salt solution (HBSS, ThermoFisher) and treated with 1 mM ethylenediaminetetraacetic acid (EDTA, J. T. Baker) under rotation for 2 × 45 minutes at 37°C to separate the epithelium from the lamina propria and release the intraepithelial lymphocytes. To obtain single cells from the lamina propria, biopsies were subsequently washed with HBSS and incubated with 5 mL Iscove’s Modified Dulbecco’s Media (IMDM) (Lonza) containing 2% Fetal Calf Serum (FCS) (Bodinco), 1,000U/mL collagenase IV (Worthington), and 10 mg/mL DNase I grade II (Roche Diagnostics) for 1.5 hours at 37°C. Cells isolated from both the epithelium and lamina propria were then pooled and filtered through a 70 μm cell strainer and collected by centrifugation. To remove dead cells, the cell pellets were incubated with 10 mg/mL DNase I in 5 ml 2% FCS/IMDM and filtered through a 30 μm cell strainer. The single-cell suspensions were collected after centrifugation and resuspended in IMDM complete medium supplemented with 2 mM L-glutamine (Lonza), 10% human serum, 100 U/mL Penicillin-Streptomycin (ThermoFisher), 0.25 ug/ml Amphotericin B (Gibco), 100 μg/mL Normocin (InvivoGen) at 4°C for future steps.

PBMCs were isolated from freshly drawn heparin anticoagulated blood samples using Ficoll-Paque™ density-gradient centrifugation. Isolated cells were washed with phosphate-buffered saline (PBS), counted, and 1 million cells were stored in 20% FCS/RPMI at 4°C for future steps.

### T-cell activation assay

Non-tissue culture-treated plates were incubated with 4 μg/mL goat anti-mouse IgG (Jackson ImmunoResearch) at 4°C overnight in PBS and subsequently with 2.5 μg/mL α-CD3 (clone Hit3a, ThermoFisher) at 37 °C for 3 hours in PBS. The fresh intestinal single cells and PBMCs were seeded in the α-CD3 pre-coated plates with 1 μg/mL α-CD28 (clone CD28.2, ThermoFisher) in complete medium for 6 h at 37 °C. In cohort 1, brefeldin A (1:1000, Sigma-Aldrich) was added for the final 5 hours; in cohort 2, monensin (1:1500, BD GolgiStop™) was added for the final 3 h. Monensin and brefeldin A were used in separate settings but yielded comparable cytokine retention efficiency. Data from both experiments were therefore combined.

### Full-spectrum flow cytometry

For surface staining, cells were incubated with fluorochrome-conjugated antibodies, human Fc block and monocyte block (BioLegend) for 30 min at 4 °C in 0.5% FCS/PBS. Cells were then fixed/permeabilized using Foxp3/Transcription Factor Staining Buffer Set (ThermoFisher). Subsequently, intracellular staining for transcription factors, cytokines and Ki67 was performed for 45 min at 4 °C in Perm Wash Buffer (ThermoFisher). Samples were measured on 5-laser Aurora (Cytek), and unmixed with single stained controls on PBMCs or UltraComp eBeads (ThermoFisher) on the SpectroFlo platform (Cytek). Unstained reference controls used to subtract autofluorescence were paired to their corresponding fully stained samples. Sample processing, staining and measurement for phenotypic identification were performed within the same day; cells for cytokine profile identification were fixed and stored at 4 °C in 0.5% FCS/PBS overnight, subsequently stained with intracellular antibodies and measured the following day. To account for technical variation, reference samples of PBMCs and tonsillar mononuclear cells from single donors were included in every spectral cytometry experiment. Data were analyzed with FlowJo software version 10 (Tree Star Inc) and OMIQ online platform (https://www.omiq.ai/, Inc., Santa Clara, CA, USA). All antibodies used are listed in Supplementary Table 2 and 3.

### Multispectral immunofluorescence and microscopy

Freshly obtained biopsies were formalin-fixed and paraffin-embedded by the Department of Pathology at LUMC. 4 μm sections were cut using a microtome, mounted on silane-coated slides, and dried overnight at 37 °C. Sections were deparaffinized using three rounds of xylene, followed by three 100% ethanol washes, blocking of endogenous peroxidase for 20 min, and then rinsed with 70% and 50% ethonal and Milli-Q water. For antigen retrieval, sections were boiled in 1x Tris-EDTA buffer (10 mM, pH 9) for 10 min and allowed to cool down for ± 60 min. Sections were stained in the dark in 5 staining cycles. Each staining cycle had the following procedure: All sections were blocked by Superblock (Thermofisher) for 10 min at room temperature (RT). Excess superblock was removed, and slides were incubated with primary antibody for 1hr at RT. After incubation, tyramide signal amplification was performed with poly-HRP (immunologic) for 10 min at RT, followed by incubation with Opal (Akoya Biosciences) for 10 min in the dark at RT according to manufacturer’s protocol. Next, sections were boiled in 1x citrate buffer (10 mM, pH 6) for 15 min to permit further staining cycles. The primary staining sequence followed the order of CD3, E-cadherin, CD103, CD8 and CD4 (antibodies used are listed in Supplementary Table 4). The opal staining sequence followed the order of 520, 690, 620, 650 and 570. Lastly, the sections were incubated with 1 μM DAPI for 15 min at RT for nuclear counterstaining, and mounted using ProLong Gold Antifade Reagent (CAT#9071, CST). Each ROI was imaged at 20× magnification with the Vectra 3.0 Automated Quantitative Pathology Imaging System (Perkin Elmer). Inform software (Akoya) was used for spectral separation of Opals. Qupath software (version 0.6.0) was further used for image analysis.

### Data analysis

Dimensionality reduction (opt-SNE) on CD3^+^ T cell data was performed on OMIQ following the R package “PeacoQC” to perform automated cleaning of anomalies and arcsinh-transformation. Subsequently, pre-processed CD4^+^ memory T cells and CD8α^+^ T cells were exported and analyzed by t-SNE (each downsampled to 1,000 cells for visualizations) in Cytosplore version 2.3.1^16^. The t-SNE computation dynamics to predict the trajectory of CD4 memory T cells was performed by Cytosplore, as demonstrated in our previous work^15^. Wishbone analysis of IBD-associated clusters was conducted in OMIQ, with CD4-13 cluster as starting point. PCA of the samples for cell cluster frequencies was performed using the R-package ‘pca3d’, where the centroid for each patient group is plotted in a scatter plot. The R package ‘corrplot’ was used to calculate the correlation networks of cell frequencies of the identified cell subsets.

### Statistics

Data were presented as median and interquartile range. Patient group comparisons were performed using two-tailed Kruskal–Wallis test followed by Dunn’s test for multiple comparisons. Cytokine proportions between the noninflamed and inflamed groups (Fig. 5D, E, and G) were analyzed using two-tailed Mann–Whitney tests, while comparisons between the HLA-DR⁻CD38⁻ and HLA-DR⁺CD38⁺ populations (Fig. 6I) were performed using Wilcoxon tests (GraphPad Prism version 10.2.3). Linear Mixed-effects model implemented in R was used to account for stimulation effects within each inflammation group (Fig. 5A–C).

## Data availability

Data will be made publicly available on Flow Repository upon publication of the manuscript.

## Acknowledgments

We thank all the patients for participating in this study. We thank all gastroenterologists and nurses for performing endoscopies and obtaining patient materials. We thank Zhixian Wang, Yizhi Wang, Yanling Xiao, Oscar R. J. van Hengel, Zheng Li and Noel F. de Miranda for valuable contributions to experimental setting and data analysis. In addition, we thank groupmates for insightful discussions and assistance with experiments, and all the operators in the Flow Cytometry Core Facility of the LUMC for their technical assistance during measurements on Cytek Aurora.

## Author contributions

QJ, FK, MFP, AM and VvU conceived the study and wrote the manuscript. AvdM-dJ, CRM-B, PV and LFO offered clinical advice and assisted in biopsies collection. QJ performed the experiments and analyzed the data. NG analyzed part of PBMC data. VAM helped to perform the staining of tissue sections, TH provided conceptual input for trajectory analysis, and CL helped to perform the isolation of single cells.

