## Supplementary materials for "Tissue-embedded CD4⁺ plasticity defines mucosal immunity in Inflammatory Bowel Disease"

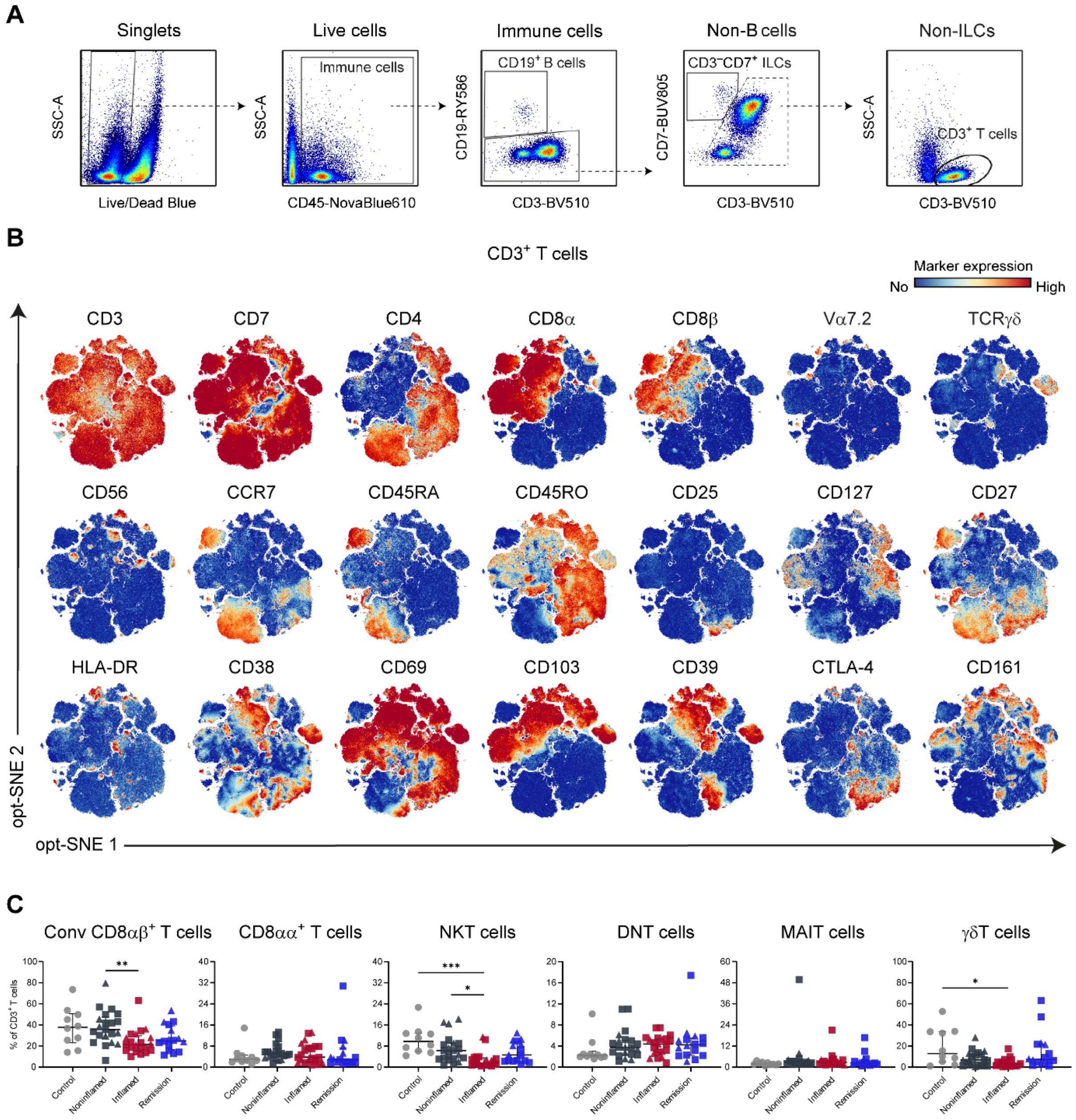


**Supplementary Fig. 1. Gating strategy and distribution of intestinal T-cell subsets across disease states. (A),** Bivariate flow cytometry plots showing the gating strategy used to exclude dead cells, $CD19^+$ B cells, $CD3^-CD7^+$ innate lymphoid cells (ILCs) and to identify intestinal $CD3^+$ T cells. **(B),** opt-SNE embedding plot showing 3.5 million live $CD45^+$ cells, colored by the expression of the indicated markers. **(C)**, Dot plots showing the frequencies of conventional (Conv) $CD8\alpha\beta^+$ T, $CD8\alpha\alpha^+$ T, NKT ($CD56^+CD8\alpha\beta^+$), Double Negative (DN, $CD8\alpha\beta^-CD4^-$), MAIT and γδ T cells, stratified by disease state. Error bars indicate median with interquartile range. *p≤0.05, **p≤0.01, ***p≤0.001, Kruskal–Wallis test with Dunn's test for multiple comparisons.

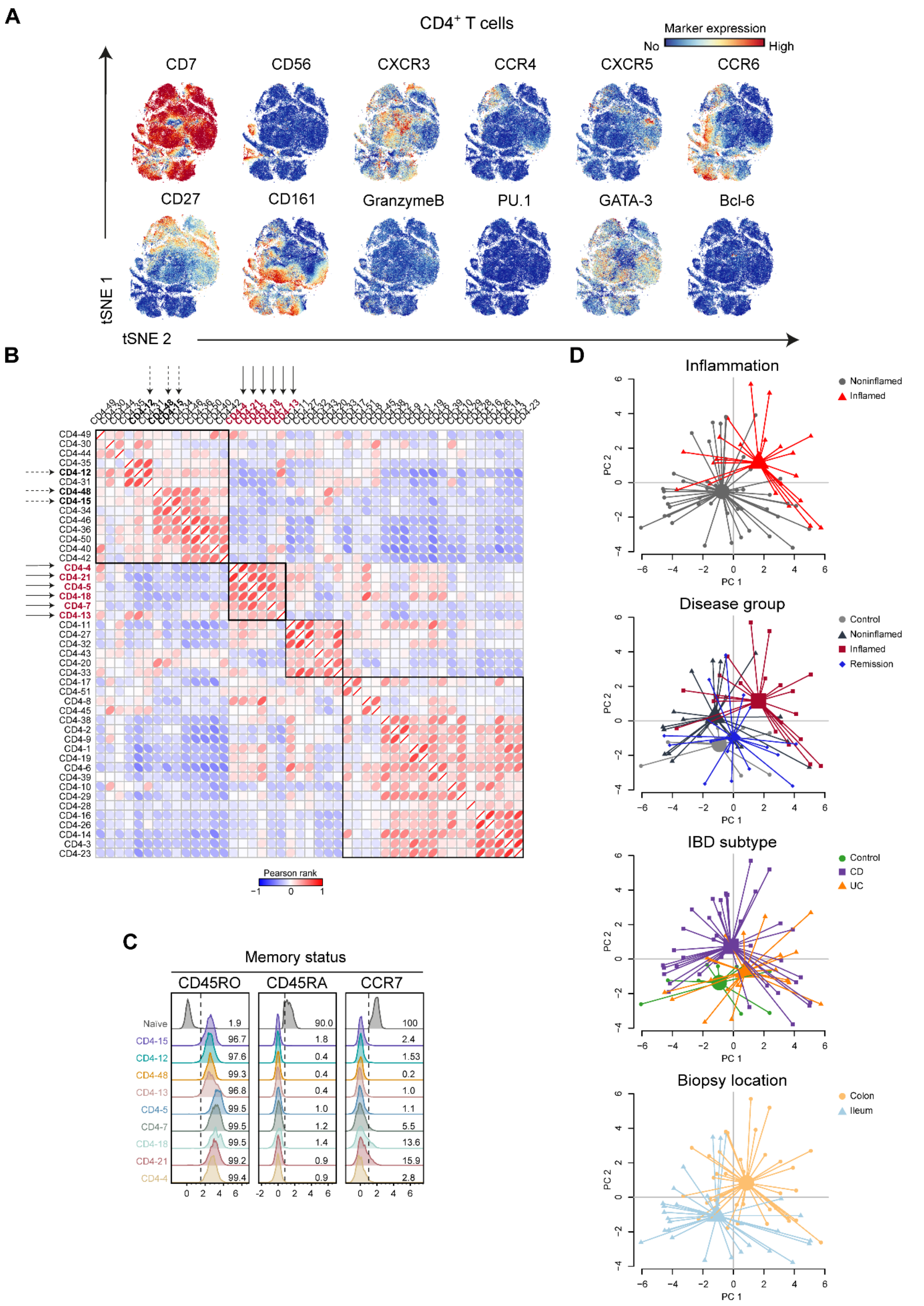


**Supplementary Fig. 2. High-dimensional phenotypic landscape and sample-level distribution of intestinal CD4+ memory T cells. (A)**, t-SNE embedding plots of 65,732 intestinal

CD4$^{+}$ memory T cells, colored by the expression values of the indicated markers. **(B)**, Pearson rank-based correlation matrix showing pairwise relationships among 46 memory CD4$^{+}$ T cell clusters based on their frequencies (% of total memory CD4$^{+}$ T cells). Hierarchical clustering of correlations delineates four major cellular network groups (black squares). Dashed arrows highlight noninflamed clusters (bold black), and solid arrows indicate inflammation-associated clusters (bold red). **(C)**, Histograms showing overlaid CD4$^{+}$ naïve T cells (random selection of 1,000 naïve cells from 68 samples) and IBD-associated clusters (derived from Fig. 2D–E), showing expression patterns indicative of memory status versus naïve status. **(D)**, PCA of 46 immune cell subsets from 68 intestinal samples (% of CD4$^{+}$ memory T cells). Each dot represents one intestinal sample, colored according to inflammation states, disease states (control, noninflamed IBD, inflamed IBD, and remission IBD), disease subgroup (control, CD and UC) and intestinal location (colon and ileum). Lines connect samples to the centroid of their respective group.

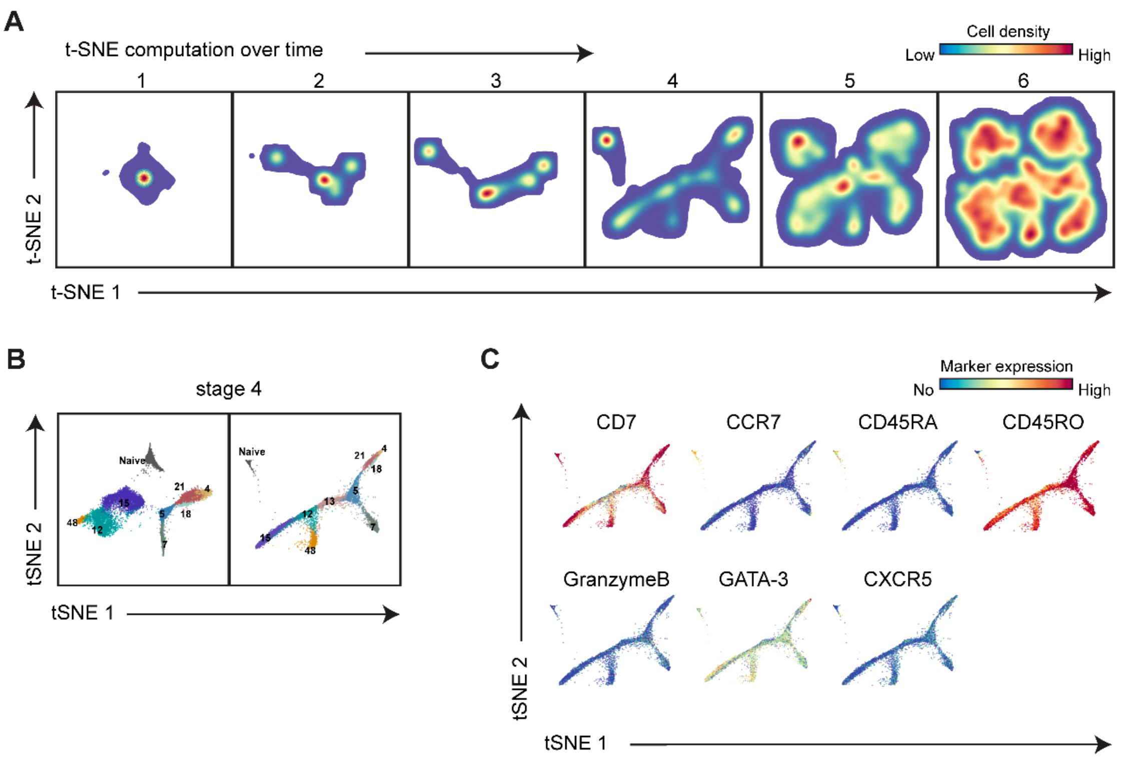


**Supplementary Fig. 3. t-SNE computation dynamics reveal structure of IBD-associated CD4$^+$ T-cell clusters. (A)**, t-SNE embeddings of CD4$^+$ naïve T cells and 9 IBD-associated CD4$^+$ T cell clusters, showing density distributions at six stages of the t-SNE computation, corresponding to Fig. 3A. **(B)**, Two t-SNE computations at stage 4, performed with (right) or without (left) inclusion of cluster CD4-13, illustrating the impact of this cluster on the embedding structure of IBD-associated clusters. **(C)**, t-SNE computation at stage 4, colored by the expression of the indicated markers, highlighting differential expression patterns in the main branches of the embedding.

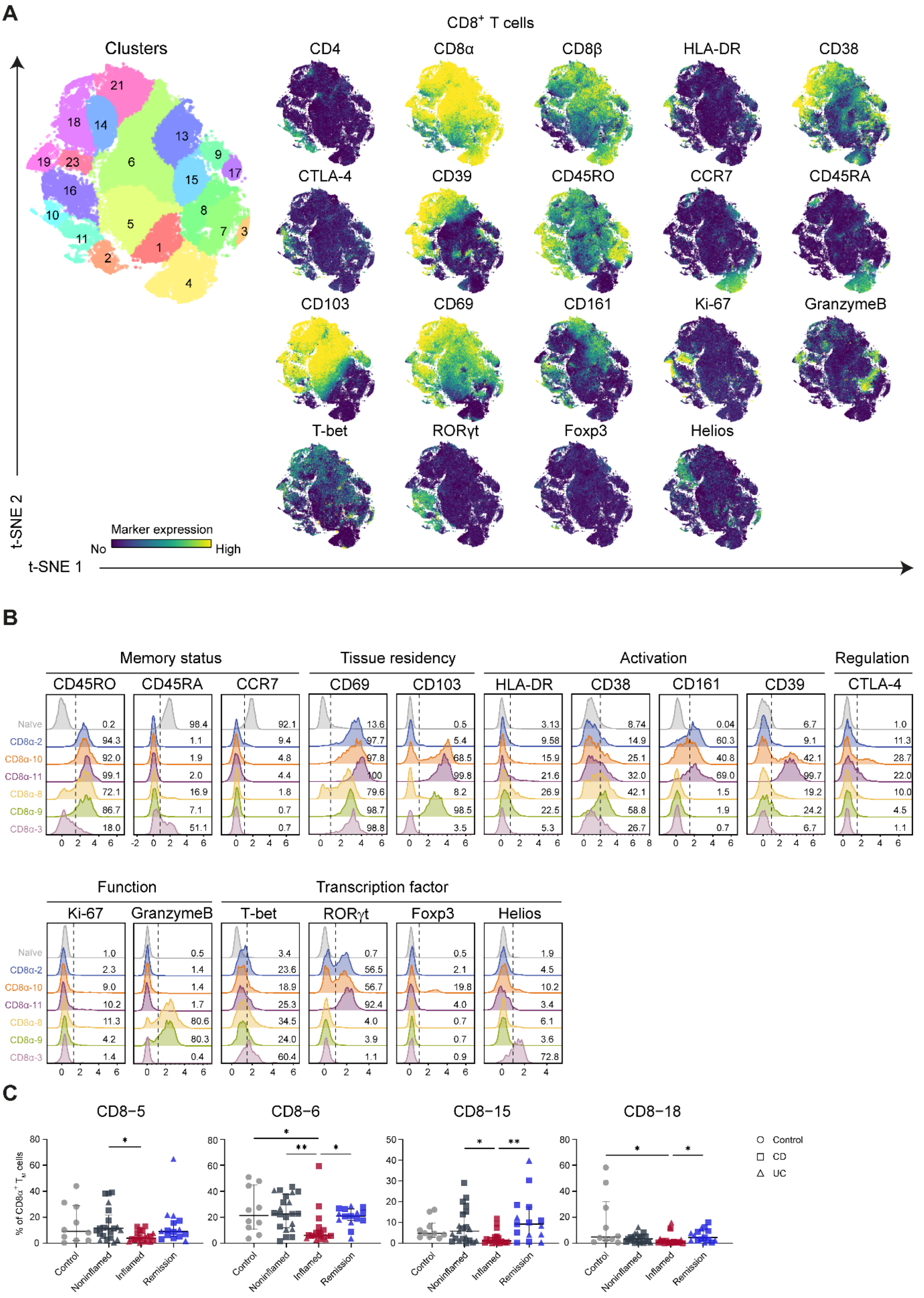


**Supplementary Fig. 4. Phenotypic landscape and inflammatory loss of intestinal CD8α⁺ T-cell subsets. (A)**, t-SNE embedding of 677,613 intestinal CD8α⁺ T cells showing cluster partitions in distinct colors (left), and the expression of the indicated markers (right). Samples were downsampled to at least 1,000 cells per sample. **(B),** Overlaid histograms of CD8α$^+$ naïve clusters and IBD-associated clusters (derived from the third network in Fig. 4A), showing expression patterns related to memory differentiation (CD45RA, CD45RO, CCR7), activation (HLA-DR, CD38 and CD39), regulation (CTLA-4), proliferation (Ki-67), cytotoxicity (Granzyme-B) and transcription factors (T-bet, RORγt, Foxp3 and Helios). **(C)**, Frequencies of CD8 T cell clusters that are reduced in inflamed tissue, stratified by disease state, expressed as percentage of total CD8α$^+$ T cells. Error bars indicate median with interquartile range. *$p \leq 0.05$, **$p \leq 0.01$, ***$p \leq 0.001$, Kruskal–Wallis test with Dunn’s test for multiple comparisons.

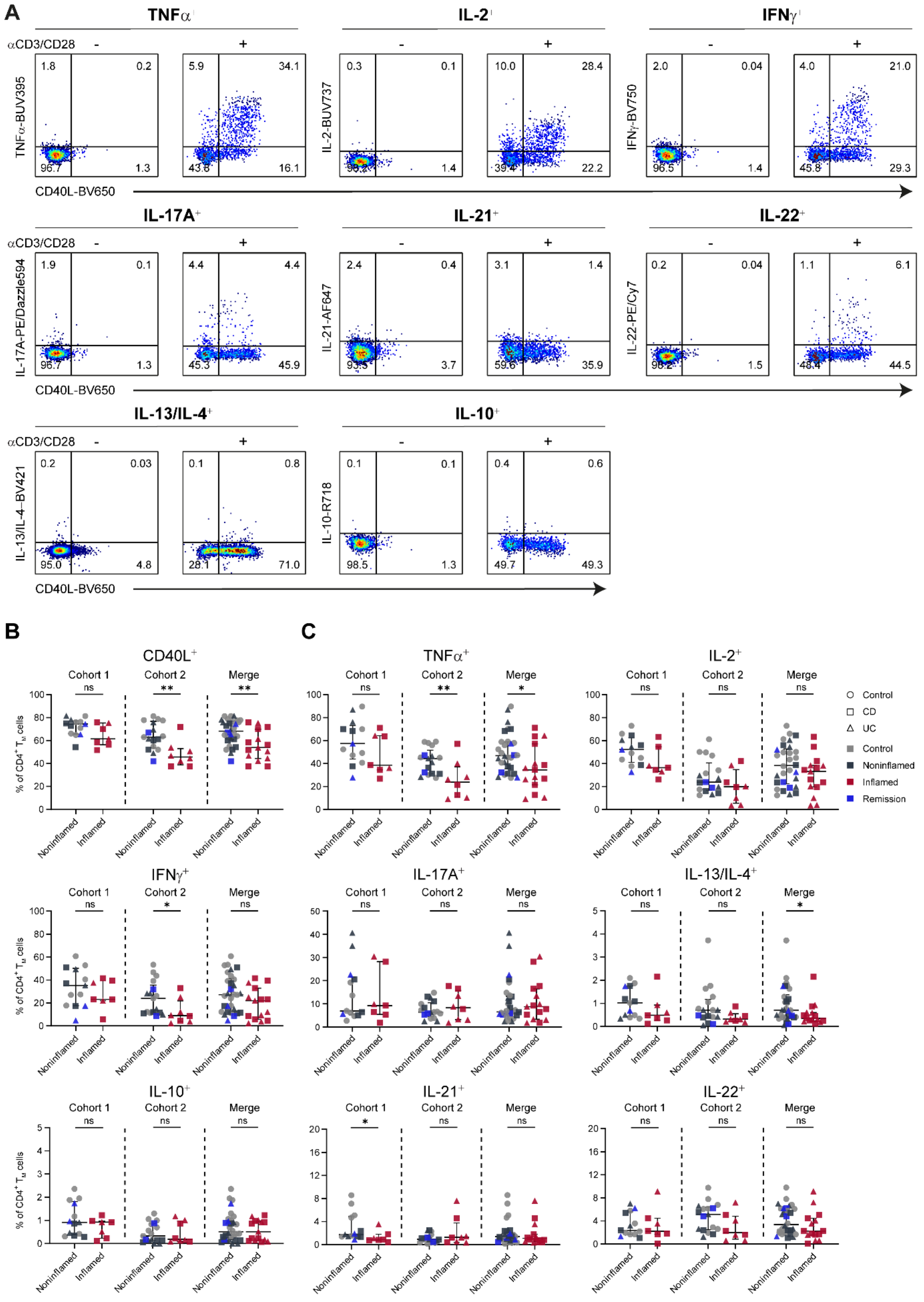


**Supplementary Fig. 5. Cytokine and CD40L responses of intestinal CD4⁺ memory T cells to TCR stimulation. (A),** Representative biaxial flow cytometry plots of intestinal CD4⁺ memory T cells after stimulation with α-CD3 and α-CD28, illustrating the gating strategy for intracellular cytokine and CD40L detection. **(B–C)**, Dot plots showing the intracellular expression of CD40L, IL-2, TNFα, IFNγ, IL-17A, IL-13/IL-4 and IL-10 in $CD4^{+}$ memory T cells from Cohort 1, Cohort 2, and the combined dataset. Error bars indicate median with interquartile range. “ns” denotes not significant, *$p \leq 0.05$, **$p \leq 0.01$, ***$p \leq 0.001$, Mann–Whitney U test.

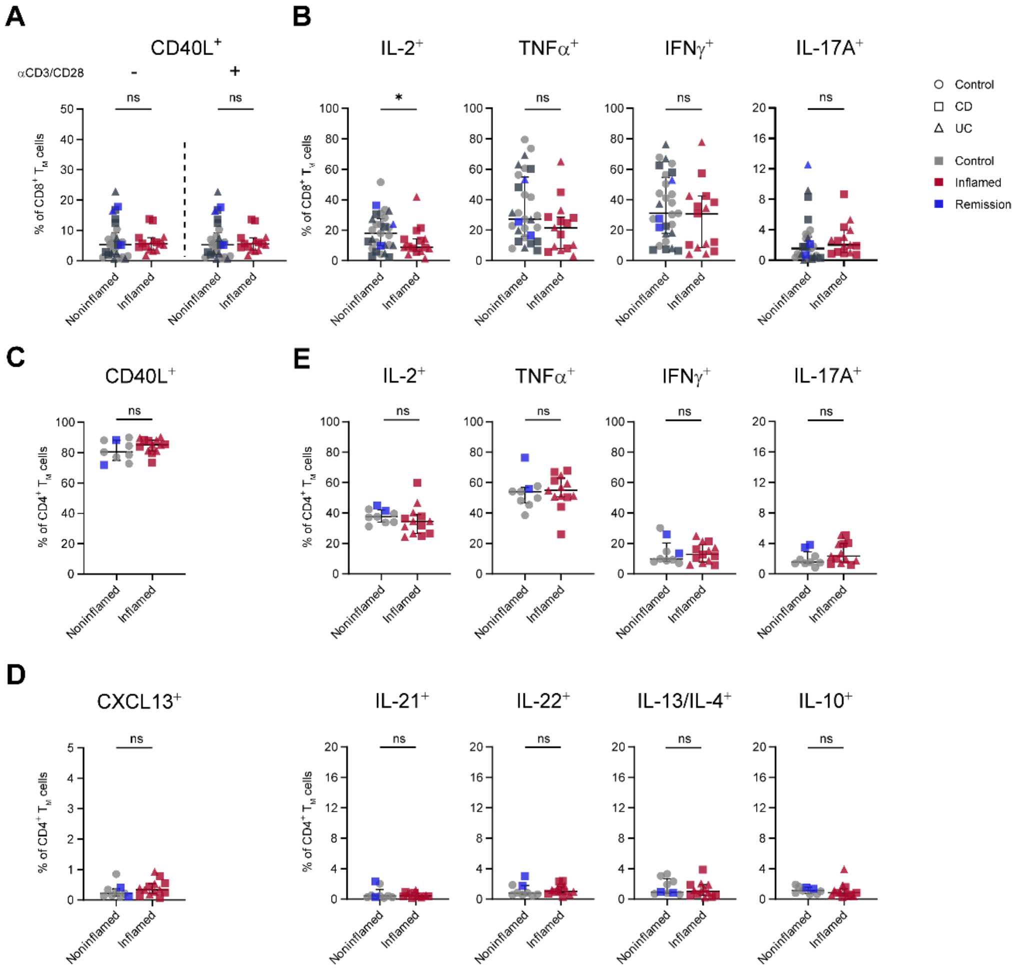


**Supplementary Fig. 6. (A–C), Cytokine responses of circulating CD4⁺ memory T cells in paired noninflamed and inflamed IBD samples.** Peripheral blood mononuclear cells from paired patients were stimulated with α-CD3/α-CD28 for 6 hours. Dot plots showing intracellular expression of CD40L, CXCL13, IL-2, TNFα, IFN-γ, IL-17A, IL-13/IL4 and IL-10 in $CD4^+$ memory T cells, comparing noninflamed and inflamed groups as determined by flow cytometry. (D-E), Cytokine responses of intestinal CD8 memory T cells upon TCR activation for 6 hours. Dot plots showing intracellular expressions of CD40L, IL-2, TNFα, IFN-γ and IL-17A. *p≤0.05, "ns" denotes not significant, Mann-Whitney U test.

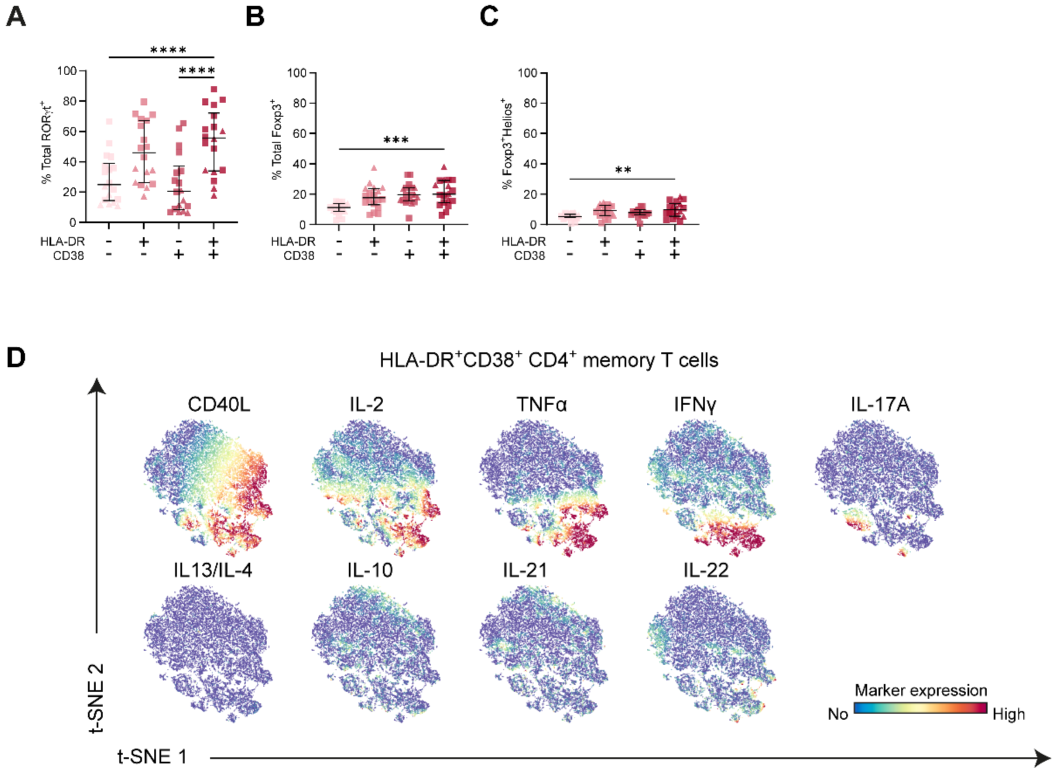


**Supplementary Fig. 7. Association of HLA-DR/CD38 activation states with RORγt$^+$ and Foxp3$^+$ CD4$^+$ T-cell subsets and cytokine responses in HLA-DR$^+$CD38$^+$ cluster. (A–C)**, Frequencies of RORγt$^+$, Foxp3$^+$ and Foxp3$^+$ Helios$^+$ CD4$^+$ T cells compared among four subsets defined by HLA-DR (positive or negative) and CD38 (positive or negative) expression. *$p \le 0.05$, **$p \le 0.01$, ***$p \le 0.001$, Kruskal–Wallis test with Dunn's post hoc test for multiple comparisons. **(D)**, t-SNE embedding of 16, 370 intestinal HLA-DR$^+$CD38$^+$ memory CD4$^+$ T cells derived from 16 inflamed samples showing the expression of the cytokines and CD40L followed by stimulation for 6 hours.

**Supplementary Table 1. Clinical metadata**

| Number | Patient | Inflammation | Disease state | Gender | Earlier inflammation | Age at endoscopy | Disease | Montreal | Severity | Location | Location (x nr. of tissues) | Peripheral blood | Treatment |
|---|---|---|---|---|---|---|---|---|---|---|---|---|---|
| **Phenotype characterization** | | | | | | | | | | | | | |
| 1 | BL52 | Noninflamed | Control | Male | | 27 | NA | NA | | Colon | 2x Unknown | Yes | |
| 2 | BL52 | Noninflamed | Control | Male | | 27 | NA | NA | | Ileum | 2x Terminal | Yes | |
| 3 | BL53 | Noninflamed | Control | Female | | 57 | NA | NA | | Colon | 2x Sigmoid | Yes | |
| 4 | BL53 | Noninflamed | Control | Female | | 57 | NA | NA | | Ileum | 2x Terminal | Yes | |
| 5 | BL54 | Noninflamed | Control | Male | | 75 | NA | NA | | Colon | 2x Descending | Yes | |
| 6 | BL54 | Noninflamed | Control | Male | | 75 | NA | NA | | Ileum | 2x Terminal | Yes | |
| 7 | A53 | Noninflamed | Control | Female | | 30 | NA | NA | | Colon | 1x Cecum+3x Ascending | Yes | |
| 8 | A56 | Noninflamed | Control | Female | | 27 | NA | NA | | Colon | 3x Transverse | Yes | |
| 9 | A53 | Noninflamed | Control | Female | | 30 | NA | NA | | Ileum | 4x Terminal | Yes | |
| 10 | A56 | Noninflamed | Control | Female | | 27 | NA | NA | | Ileum | 2x Terminal | Yes | |
| 11 | A49 | Noninflamed | Active | Male | No | 25 | CD | A2 L2L4 B1 | | Ileum | 2x Terminal | Yes | Adalimumab |
| 12 | A50 | Noninflamed | Active | Female | No | 33 | CD | A2 L2 P | | Ileum | 4x Terminal | Yes | Adalimumab and Vedolizumab |
| 13 | A51 | Noninflamed | Active | Male | Yes | 39 | CD | A2 L3 B2 | | Ileum | 2x Terminal | Yes | Adalimumab |
| 14 | A51 | Noninflamed | Active | Male | Yes | 39 | CD | A2 L3 B2 | | Colon | 4x Descending | Yes | Adalimumab |
| 15 | A52 | Noninflamed | Active | Female | Yes | 40 | CD | A2 L3 B2 | | Colon | 4x Transverse | Yes | Ustekinumab |
| 16 | A62 | Noninflamed | Active | Female | No | 39 | CD | A2 L3 B1 | | Ileum | 2x Terminal | Yes | Adalimumab |
| 17 | A67 | Noninflamed | Active | Male | No | 26 | CD | A2 L3 B2 | | Colon | 2x Sigmoid | Yes | Vedolizumab |
| 18 | A67 | Noninflamed | Active | Male | No | 26 | CD | A2 L3 B2 | | Ileum | 3x Terminal | Yes | Vedolizumab |
| 19 | BL59 | Noninflamed | Active | Male | No | 71 | UC | A3 E3 | | Ileum | 2x Terminal | Yes | 5ASA |
| 20 | BL63 | Noninflamed | Active | Male | No | 35 | UC | A1 E3 | | Ileum | 2x Terminal | Yes | 5ASA |
| 21 | BL64 | Noninflamed | Active | Female | Yes | 63 | CD | A2 L3 B2 | | Ileum | 2x Terminal | Yes | Ustekinumab |
| 22 | BL65 | Noninflamed | Active | Male | Yes | 32 | UC | A2 E3 | | Colon | 2x Descending | Yes | Ustekinumab |
| 23 | BL66 | Noninflamed | Active | Female | No | 44 | UC | A2 E2 | | Colon | 2x Ascending | Yes | Thioguanine |
| 24 | KK28 | Noninflamed | Active | Male | No | 16 | UC | A1 E3 | | Ileum | 3x Terminal | Yes | No (naive) |
| 25 | A59 | Noninflamed | Active | Female | Yes | 41 | CD | A1 L3 B2p | | Colon | 3x Ascending | Yes | Infliximab |
| 26 | A55 | Noninflamed | Active | Female | Yes | 40 | UC | A2 E3 | | Colon | 2x Ascending+1x Transverse | Yes | 5ASA |
| 27 | A60 | Noninflamed | Active | Male | Yes | 65 | CD | A2 L3 B1 | | Colon | 4x Sigmoid | Yes | Adalimumab |
| 28 | A57 | Noninflamed | Active | Male | Yes | 36 | CD | A1 L3 | | Ileum | 3x Terminal | Yes | Adalimumab |
| 29 | A65 | Noninflamed | Active | Female | Yes | 23 | CD | A1 L3 B1 | | Ileum | 2x Terminal | Yes | Infliximab |
| 30 | KK26 | Noninflamed | Active | Female | Yes | 15 | CD | A1 L3 B1 | | Ileum | 3x Terminal | Yes | 5ASA |
| 31 | A55 | Noninflamed | Active | Female | Yes | 40 | UC | A2 E3 | | Colon | 1x Rectum | Yes | 5ASA |
| 32 | A49 | Inflamed | Active | Male | | 25 | CD | A2 L2L4 B1 | Moderate | Colon | 2x Ascending+2x Descending | Yes | Adalimumab |
| 33 | A50 | Inflamed | Active | Female | | 33 | CD | A2 L2 P | Severe | Colon | 4x Descending | Yes | Adalimumab and Vedolizumab |
| 34 | A51 | Inflamed | Active | Male | | 39 | CD | A2 L3 B2 | Mild | Ileum | 2x Terminal | Yes | Adalimumab |
| 35 | A52 | Inflamed | Active | Female | | 40 | CD | A2 L3 B2 | Severe | Colon | 4x Descending | Yes | Ustekinumab |
| 36 | A61 | Inflamed | Active | Male | | 31 | UC | A2 E3 | Moderate | Colon | 4x Ascending+3x Cecum | Yes | Budesonide |
| 37 | A62 | Inflamed | Active | Female | | 39 | CD | A2 L3 B1 | Mild | Colon | 2x Transverse | Yes | Adalimumab |
| 38 | A62 | Inflamed | Active | Female | | 39 | CD | A2 L3 B1 | Mild | Ileum | 2x Terminal | Yes | Adalimumab |
| 39 | A67 | Inflamed | Active | Male | | 26 | CD | A2 L3 B2 | Mild | Colon | 3x Cecum | Yes | Vedolizumab |
| 40 | BL59 | Inflamed | Active | Male | | 71 | UC | A3 E3 | Mild | Colon | 2x Sigmoid | Yes | 5ASA |
| 41 | BL63 | Inflamed | Active | Male | | 35 | UC | A1 E3 | Mild | Colon | 2x Ascending | Yes | 5ASA |
| 42 | BL64 | Inflamed | Active | Female | | 63 | CD | A2 L3 B2 | Mild | Ileum | 2x Terminal | Yes | Ustekinumab |
| 43 | BL65 | Inflamed | Active | Male | | 32 | UC | A2 E3 | Mild | Colon | 2x Rectum | Yes | Ustekinumab |
| 44 | BL66 | Inflamed | Active | Female | | 44 | UC | A2 E2 | Mild | Colon | 2x Sigmoid | Yes | Thioguanine |
| 45 | A65 | Inflamed | Active | Female | | 23 | CD | A1 L3 B1 | Mild | Colon | 1x Cecum+1x Ascending+1x Transverse+1x Sigmoid | Yes | Infliximab |
| 46 | A55 | Inflamed | Active | Female | | 40 | UC | A2 E3 | Mild | Colon | 2x Cecum | Yes | 5ASA |
| 47 | A57 | Inflamed | Active | Male | | 36 | CD | A1 L3 | Mild | Colon | 3x Ascending | Yes | Adalimumab |
| 48 | A65 | Inflamed | Active | Female | | 23 | CD | A1 L3 B1 | Mild | Ileum | 2x Terminal | Yes | Infliximab |
| 49 | KK26 | Inflamed | Active | Female | | 15 | CD | A1 L3 B1 | Moderate | Ileum | 3x Terminal | Yes | 5ASA |
| 50 | A59 | Inflamed | Active | Female | | 41 | CD | A1 L3 B2p | Mild | Colon | 3x Sigmoid | Yes | Infliximab |
| 51 | A60 | Inflamed | Active | Male | | 65 | CD | A2 L3 B1 | Mild | Colon | 4x Transverse | Yes | Adalimumab |
| 52 | KK28 | Inflamed | Active | Male | | 16 | UC | A1 E3 | Severe | Colon | 1x Ascending+2x Descending | Yes | No (naive) |
| 53 | A68 | Noninflamed | Remission | Male | Yes | 21 | UC | A1 E3 | | Colon | 3x Ascending | Yes | No medication |
| 54 | A68 | Noninflamed | Remission | Male | No | 21 | UC | A1 E3 | | Ileum | 3x Terminal | Yes | No medication |
| 55 | A73 | Noninflamed | Remission | Male | Yes | 23 | UC | A1 E3 | | Colon | 4x Descending | Yes | Vedolizumab |
| 56 | A73 | Noninflamed | Remission | Male | No | 23 | UC | A1 E3 | | Ileum | 4x Terminal | Yes | Vedolizumab |
| 57 | BL44 | Noninflamed | Remission | Male | Yes | 30 | CD | A2 L3 B1 | | Colon | 4x Unknown | Yes | 5ASA |
| 58 | BL44 | Noninflamed | Remission | Male | Yes | 30 | CD | A2 L3 B1 | | Ileum | 4x Terminal | Yes | 5ASA |
| 59 | BL51 | Noninflamed | Remission | Female | Yes | 63 | CD | A2 L3 B2 | | Colon | 4x Unknown | Yes | Adalimumab |
| 60 | BL51 | Noninflamed | Remission | Female | Yes | 63 | CD | A2 L3 B2 | | Ileum | 4x Terminal | Yes | Adalimumab |
| 61 | BL60 | Noninflamed | Remission | Male | Yes | 25 | CD | A1 L2 B1 | | Colon | 2x Unknown | Yes | Adalimumab |
| 62 | BL60 | Noninflamed | Remission | Male | No | 25 | CD | A1 L2 B1 | | Ileum | 2x Terminal | Yes | Adalimumab |
| 63 | A63 | Noninflamed | Remission | Female | No | 30 | CD | A1 L1L4 B1p | | Colon | 2x Ascending+2x Sigmoid | Yes | Infliximab |
| 64 | A64 | Noninflamed | Remission | Male | No | 33 | UC | A2 E1 | | Colon | 2x Ascending+2x Sigmoid+2x Transverse | Yes | Infliximab |
| 65 | A66 | Noninflamed | Remission | Female | No | 33 | CD | A2 L1 B2 | | Colon | 4x Ascending | Yes | Thioguanine |
| 66 | A63 | Noninflamed | Remission | Female | Yes | 30 | CD | A1 L1L4 B1p | | Ileum | 2x Terminal | Yes | Infliximab |
| 67 | A64 | Noninflamed | Remission | Male | No | 33 | UC | A2 E1 | | Ileum | 2x Terminal | Yes | Infliximab |
| 68 | A66 | Noninflamed | Remission | Female | Yes | 33 | CD | A2 L1 B2 | | Ileum | 4x Terminal | Yes | Thioguanie |
| **TCR stimulation_First Cohort** | | | | | | | | | | | | | |
| 69 | KK30 | Noninflamed | Control | Female | | 13 | NA | NA | | Ileum | 2x Unknown | Yes | |
| 70 | KK30 | Noninflamed | Control | Female | | 13 | NA | NA | | Ileum | 2x Unknown | Yes | |
| 71 | BL46 | Noninflamed | Control | Female | | 69 | NA | NA | | Colon | 4x Unknown | Yes | |
| 72 | BL46 | Noninflamed | Control | Female | | 69 | NA | NA | | Ileum | 4x Terminal | Yes | |
| 73 | BL47 | Noninflamed | Control | Female | | 46 | NA | NA | | Colon | 4x Unknown | Yes | |
| 74 | BL47 | Noninflamed | Control | Female | | 46 | NA | NA | | Ileum | 4x Terminal | Yes | |
| 75 | A71 | Noninflamed | Active | Female | No | 51 | CD | A2 L2 B1 | | Ileum | 2x Terminal | Yes | Ustekinumab |
| 76 | A72 | Noninflamed | Active | Female | Yes | 45 | CD | A3 L2 B1 | | Colon | 2x Sigmoid | Yes | Vedolizumab |
| 77 | BL45 | Noninflamed | Active | Female | Yes | 49 | UC | A1 E3 | | Colon | 3x Ascending | Yes | Tofacitinib |
| 78 | BL45 | Noninflamed | Active | Female | No | 49 | UC | A1 E3 | | Ileum | 2x Terminal | Yes | Tofacitinib |
| 79 | A75 | Noninflamed | Active | Female | Yes | 47 | CD | A2 L3 B2p | | Colon | 3x Transverse | Yes | Ustekinumab |
| 80 | A71 | Inflamed | Active | Female | | 51 | CD | A2 L2 B1 | Moderate | Colon | 4x Ascending | Yes | Ustekinumab |
| 81 | A71 | Inflamed | Active | Female | | 51 | CD | A2 L2 B1 | Mild | Ileum | 1x Terminal | Yes | Ustekinumab |
| 82 | A72 | Inflamed | Active | Female | | 45 | CD | A3 L2 B1 | Moderate | Colon | 2x Ascending+2x Transverse | Yes | Vedolizumab |
| 83 | A72 | Inflamed | Active | Female | | 45 | CD | A3 L2 B1 | Moderate | Colon | 4x Cecum | Yes | Vedolizumab |
| 84 | BL45 | Inflamed | Active | Female | | 49 | UC | A1 E3 | Mild | Colon | 2x Sigmoid | Yes | Tofacitinib |
| 85 | A75 | Inflamed | Active | Female | | 47 | CD | A2 L3 B2p | Moderate | Ileum | 3x Terminal | Yes | Ustekinumab |
| 86 | KK29 | Inflamed | Active | Female | | 12 | UC | A1 E3 | Mild | Colon | 3x Descending+1x Ascending | Yes | No (naive) |
| 87 | A74 | Noninflamed | Remission | Male | Yes | 44 | UC | A2 E3 | | Colon | 4x Ascending | No | Tofacitinib, prednison 5 mg |
| 88 | A74 | Noninflamed | Remission | Male | Yes | 44 | UC | A2 E3 | | Colon | 3x Sigmoid | No | Tofacitinib, prednison 5 mg |
| **TCR stimulation_Second Cohort** | | | | | | | | | | | | | |
| 89 | BL52 | Noninflamed | Control | Male | | 27 | NA | NA | | Colon | 4x Unknown | Yes | |
| 90 | BL52 | Noninflamed | Control | Male | | 27 | NA | NA | | Ileum | 4x Terminal | Yes | |
| 91 | BL53 | Noninflamed | Control | Female | | 57 | NA | NA | | Colon | 4x Sigmoid | Yes | |
| 92 | BL53 | Noninflamed | Control | Female | | 57 | NA | NA | | Ileum | 4x Terminal | Yes | |
| 93 | BL54 | Noninflamed | Control | Male | | 75 | NA | NA | | Colon | 4x Descending | Yes | |
| 94 | BL54 | Noninflamed | Control | Male | | 75 | NA | NA | | Ileum | 4x Terminal | Yes | |
| 95 | BL55 | Noninflamed | Control | Male | | 60 | NA | NA | | Colon | 4x Unknown | No | |
| 96 | BL55 | Noninflamed | Control | Male | | 60 | NA | NA | | Ileum | 4x Terminal | No | |
| 97 | BL62 | Noninflamed | Active | Male | Yes | 67 | CD | A2 L3 | | Ileum | 4x Terminal | No | Ustekinumab |
| 98 | BL62 | Noninflamed | Active | Male | No | 67 | CD | A2 L3 | | Colon | 4x Unknown | No | Ustekinumab |
| 99 | BL64 | Noninflamed | Active | Female | Yes | 63 | CD | A2 L3 B2 | | Ileum | 4x Terminal | Yes | Ustekinumab |
| 100 | BL59 | Noninflamed | Active | Male | No | 71 | UC | A3 E3 | | Ileum | 4x Terminal | Yes | 5ASA |
| 101 | BL63 | Noninflamed | Active | Male | No | 35 | UC | A1 E3 | | Ileum | 4x Terminal | Yes | 5ASA |
| 102 | BL65 | Noninflamed | Active | Male | Yes | 32 | UC | A2 E3 | | Colon | 4x Descending | Yes | Ustekinumab |
| 103 | BL66 | Noninflamed | Active | Female | No | 44 | UC | A2 E2 | | Colon | 4x Ascending | Yes | Thioguanine |
| 104 | BL58 | Inflamed | Active | Female | | 49 | CD | A2 L1 | Mild | Ileum | 4x Terminal | No | Vedolizumab |
| 105 | BL62 | Inflamed | Active | Male | | 67 | CD | A2 L3 | Severe | Ileum | 4x Terminal | No | Ustekinumab |
| 106 | BL64 | Inflamed | Active | Female | | 63 | CD | A2 L3 B2 | Mild | Ileum | 4x Terminal | Yes | Ustekinumab |
| 107 | BL59 | Inflamed | Active | Male | | 71 | UC | A3 E3 | Mild | Colon | 4x Sigmoid | Yes | 5ASA |
| 108 | BL61 | Inflamed | Active | Female | | 38 | UC | A2 E3 | Moderate | Colon | 4x Rectum | No | 5ASA |
| 109 | BL63 | Inflamed | Active | Male | | 35 | UC | A1 E3 | Mild | Colon | 4x Ascending | Yes | 5ASA |
| 110 | BL65 | Inflamed | Active | Male | | 32 | UC | A2 E3 | Mild | Colon | 4x Rectum | Yes | Ustekinumab |
| 111 | BL66 | Inflamed | Active | Female | | 44 | UC | A2 E2 | Mild | Colon | 4x Sigmoid | Yes | Thioguanine |
| 112 | BL51 | Noninflamed | Remission | Female | Yes | 63 | CD | A2 L3 B2 | | Colon | 2x Terminal | Yes | Adalimumab |
| 113 | BL60 | Noninflamed | Remission | Male | No | 25 | CD | A1 L2 B1 | | Ileum | 4x Terminal | Yes | Adalimumab |

**Supplementary Table 2. Antibodies used in the phenotypic characterization cohort.**

| Specificity | Fluorochrome | Clone | Vendor | Catalog # |
|---|---|---|---|---|
| CD45RA | PerCP | HI100 | Biolegend | 304155 |
| CD45RO | BV570 | UCHL1 | Biolegend | 304225 |
| Viability | LIVE/DEAD™ Blue | - | Invitrogen | L34961 |
| CD25 | BV421 | BC96 | Biolegend | 302630 |
| CD27 | FITC | M-T271 | BD | 555440 |
| CD38 | APC/Fire 810 | HIT2 | Biolegend | 303550 |
| CD39 | BV650 | TU66 | BD | 563681 |
| CD69 | BUV737 | FN50 | BD | 612817 |
| CD103 | PE/Fire 700 | Ber-ACT8 | Biolegend | 350240 |
| CD127 | BV711 | A019D5 | Biolegend | 351328 |
| CD161 | BUV563 | HP-3G10 | BD | 749223 |
| CD183 (CXCR3) | APC/Fire 750 | G025H7 | Biolegend | 353754 |
| CD185 (CXCR5) | BV750 | J252D4 | Biolegend | 356942 |
| CD194 (CCR4) | BUV615 | 1G1 | BD | 613000 |
| CD196 (CCR6) | BB700 | 11A9 | BD | 566477 |
| CD197 (CCR7) | Spark NIR 685 | G043H7 | Biolegend | 353258 |
| HLA-DR | BV480 | G46-6 | BD | 566154 |
| CD3 | BV510 | UCHT1 | Biolegend | 300448 |
| CD4 | Alexa Fluor 532 | RPA-T4 | Invitrogen | 58-0049-42 |
| CD7 | BUV805 | M-T701 | BD | 742002 |
| CD8α | Spark Blue 550 | SK1 | Biolegend | 344759 |
| CD8β | BUV496 | 2ST8.5H7 | BD | 749837 |
| CD19 | Real Yellow 586 | SJ25C1 | BD | 568107 |
| CD45 | NovaFluor Blue 610-30S | 2D1 | Invitrogen | H005T03B05 |
| CD56 | Super Bright 436 | TULY56 | Invitrogen | 62-0566-41 |
| TCR Vα7.2 | BV785 | 3C10 | Biolegend | 351722 |
| TCR γδ | VioBlue | 11F2 | Miltenyi Biotec | 130-113-507 |
| Bcl-6 | R718 | K112-91 | BD | 566979 |
| CD152 (CTLA-4) | BV605 | BNI3 | Biolegend | 369610 |
| FoxP3 | APC | PCH101 | Invitrogen | 17-4776-42 |
| GATA3 | PE/Cy5 | TWAJ | Invitrogen | 15-9966-42 |
| Granzyme-B | PE-Dazzle 594 | QA16A02 | Biolegend | 372216 |
| Helios | PerCP-eFluor 710 | 22F6 | Invitrogen | 46-9883-42 |
| Ki-67 | BUV395 | B56 | BD | 564071 |
| PU.1 (SPI1) | Alexa Fluor 647 | 7C6B05 | Biolegend | 658004 |
| RORγt | PE | Q21-559 | BD | 563081 |
| T-bet | PE/Cy7 | eBio4B10 | Invitrogen | 25-5825-80 |

**Supplementary Table 3. Antibodies used in the T-cell receptor stimulation cohort.**

| Specificity | Fluorochrome | Clone | Vendor | Catalog # |
|---|---|---|---|---|
| CD45RA | PerCP | HI100 | Biolegend | 304155 |
| CD45RO | BV570 | UCHL1 | Biolegend | 304225 |
| Viability | LIVE/DEAD™ Blue | - | Invitrogen | L34961 |
| CD27 | FITC | M-T271 | BD | 555440 |
| CD38 | APC/Fire 810 | HIT2 | Biolegend | 303550 |
| CD39 | BV650 | TU66 | BD | 563681 |
| CD103 | PE/Fire 700 | Ber-ACT8 | Biolegend | 350240 |
| CD161 | BUV563 | HP-3G10 | BD | 749223 |
| CD194 (CCR4) | BUV615 | 1G1 | BD | 613000 |
| CD196 (CCR6) | BB700 | 11A9 | BD | 566477 |
| CD197 (CCR7) | Spark NIR 685 | G043H7 | Biolegend | 353258 |
| HLA-DR | BV480 | G46-6 | BD | 566154 |
| CD4 | Alexa Fluor 532 | RPA-T4 | Invitrogen | 58-0049-42 |
| CD7 | BUV805 | M-T701 | BD | 742002 |
| CD8α | Spark Blue 550 | SK1 | Biolegend | 344759 |
| CD8β | BUV496 | 2ST8.5H7 | BD | 749837 |
| CD19 | Real Yellow 586 | SJ25C1 | BD | 568107 |
| CD45 | NovaFluor Blue 610-30S | 2D1 | Invitrogen | H005T03B05 |
| CD56 | Super Bright 436 | TULY56 | Invitrogen | 62-0566-41 |
| CD3 | BV510 | UCHT1 | Biolegend | 300448 |
| TCR Vα7.2 | BV785 | 3C10 | Biolegend | 351722 |
| TCR γδ | VioBlue | 11F2 | Miltenyi Biotec | 130-113-507 |
| CD74 | BV711 | LN2 | BD | 743735 |
| CD154 (CD40L) | BV605 | 24-31 | Biolegend | 310826 |
| FoxP3 | APC | PCH101 | Invitrogen | 17-4776-42 |
| Helios | PerCP-eFluor 710 | 22F6 | Invitrogen | 46-9883-42 |
| IFNγ | BV750 | B27 | BD | 566357 |
| TNFα | BUV395 | MAb11 | BD | 563996 |
| IL-2 | BUV737 | mq1-17h12 | BD | 612836 |
| IL-10 | R718 | JES3-19F1 | BD | 567059 |
| IL-4 | BV421 | MP4-25D2 | Biolegend | 500826 |
| IL-13 | BV421 | JES10-5A2 | Biolegend | 501916 |
| IL-17A | PE-Dazzle 594 | BL168 | Biolegend | 512335 |
| IL-21 | Alexa Fluor 647 | 3A3-N2 | BD | 560493 |
| IL-22 | PE/Cy7 | 22URTI | Invitrogen | 15518646 |
| CXCL13 | PE | 53610 | Invitrogen | MA5-23832 |

**Supplementary Table 4. Antibodies used for multispectral immunofluorescence.**

| Specificity | Clone | Vendor | Catalog # | Opal |
|---|---|---|---|---|
| CD3 | D7A6E | CST | 85061BF | 520 |
| E-cadherin | HECD-1 | Biomimiq | - | 690 |
| CD103 | EP206 | CST | 95835S | 620 |
| CD8 | D8A8Y | CST | 85336S | 650 |
| CD4 | EPR6855 | Abcam | ab133616 | 570 |
